# Characterisation and reprogramming of bacteriophage mv4 integrase recombination specificity

**DOI:** 10.1101/2023.11.06.565760

**Authors:** Kevin Debatisse, Pierre Lopez, Maryse Poli, Philippe Rousseau, Manuel Campos, Michèle Coddeville, Muriel Cocaign-Bousquet, Pascal Le Bourgeois

## Abstract

Bacteriophage mv4 is a temperate bacterial virus able to integrate its genome at the 3’ end of the tRNA^SER^ of *Lactobacillus delbrueckii* subsp. *bulgaricus* chromosome through site-specific recombination. Previous investigations revealed that the ^mv4^Int/*attP*/*attB* recombination module was atypical compared to conventional heterobivalent tyrosine recombinases, such as the paradigmatic *Lambdavirus lambda* integrase, suggesting alternative recombination mechanism. *In vitro* recombination assays with random DNA libraries were used to comprehensively delineate the mv4 recombination system. We showed that ^mv4^Int is a 369-aa protein that exhibits all structural hallmarks of integrases from the Tn*916* family and interacts cooperatively with its recombination sites. We established that ^mv4^Int distinguishes itself from classical heterobivalent integrases by a greater tolerance to nucleotide variations in *attB* and core-*attP* sites. We demonstrated that, upon considering nucleotide degeneracy, the 21-bp core-*attP* and *attB* recombination sites share structural similarities with classical heterobivalent integrase systems, with two 7-bp inverted-repeat regions corresponding to ^mv4^Int core-binding sites surrounding a 7-bp strand-exchange region. Furthermore, our study highlighted compositional biases and nucleotide interdependencies within the core-binding regions that exerted a significant influence on the outcomes of recombination events.

## INTRODUCTION

Temperate bacteriophages are obligate parasites of bacterial cells that can mediate two distinct lifecycles, the lytic and lysogenic cycles. During infection, these phages typically initiate the lytic cycle, wherein the viruses hijack the host-cell machinery to replicate their DNA, assemble viral particles, and disseminate themselves in the environment by lysing the host. However, depending on several factors such as nutritional state of the host cell (1), cells or phage densities (2, 3), or small-molecule communication (4), temperate phages have the ability to enter a lysogenic cycle, which involves the repression of phage’s genes expression and the integration of the viral genome at a specific site within the host chromosome. This site-specific recombination involves site-specific recombinases (SSR) that promote DNA rearrangements between two specific DNA target sites (5). Once integrated, the bacteriophage genome, referred to as a prophage, is passively replicated alongside with the bacterial chromosome. Prophages are present in nearly half of bacterial genomes (6) and have a significant impact on bacterial genome dynamics (7, 8), accounting for up to 20 % of the bacterial DNA (9). On some occasions, such as exposure to chemicals or accumulation of DNA damages, prophage DNA excises from the bacterial chromosome and reactivates its lytic cycle (10). The integration and excision of phage DNA are mediated by a phage-encoded protein known as integrase, which primarily belongs to the heterobivalent tyrosine recombinases (YR) subfamily, although an increasing number of integrases belong to the serine integrases (11), a phylogenetically and mechanistically unrelated SSR family (5). All integrases catalyse unidirectional recombination between two specific sites, one located on the phage DNA (*attP*) and the other located on the host bacterial chromosome (*attB*). This recombination results in the integration of the prophage flanked by hybrid recombination sites (*attL* and *attR*). The reverse reaction (excision), which leads to the prophage excision by recombination between the *attL* and *attR* sites, restore the initial *attB* and *attP* sites and requires an additional phage-encoded protein, called excisionase (Xis) or recombination directionality factor (RDF), that interact with the integrase (12).

Studied for over 50 years (13), the recombination system from the coliphage *Lambdavirus lambda* represents the founding member of the heterobivalent YR subfamily. Its integrase consists of three DNA-binding domains (Figure 1A): the N-terminal arm-binding (AB) (14, 15), the core-binding (CB) (14, 16), and the C-terminal catalytic (CAT) domain containing the seven catalytic residues (R-D/E-K-H-R-H/W-Y, Figure 1B) conserved among the YRs (17). The *attP* recombination site spans approximately 240 base pairs (bp) and exhibits a complex organisation (Figure 1C) : it contains fourteen protein binding sites, including three Xis binding-sites, three IHF binding sites, one Fis binding site, and five ^λ^Int arm-binding sites (P1, P2, P’1, P’2, P’3), as well as two core-binding sites (C and C’) arranged as inverted repeats that flank the 7-bp overlap (O) sequence, the region where the strand exchange occurs (13). The *Escherichia coli attB* site is smaller (21 bp) and consists only of two inverted repeats core-binding sites (B and B’) surrounding the overlap sequence (O) identical to that of *attP* site (13). The ^λ^Int-mediated recombination reaction involves the formation of a nucleoprotein complex on *attP* called the intasome (18), wherein *attP* supercoiling is required (19), with the host factor IHF playing a fundamental architectural role (20). Once formed, this intasome can capture its cognate *attB* site (21). Extensive biochemical, functional, and structural data have facilitated the development of precise models for integrative and excisive λ recombination pathways (22–24).

**Figure 1.**
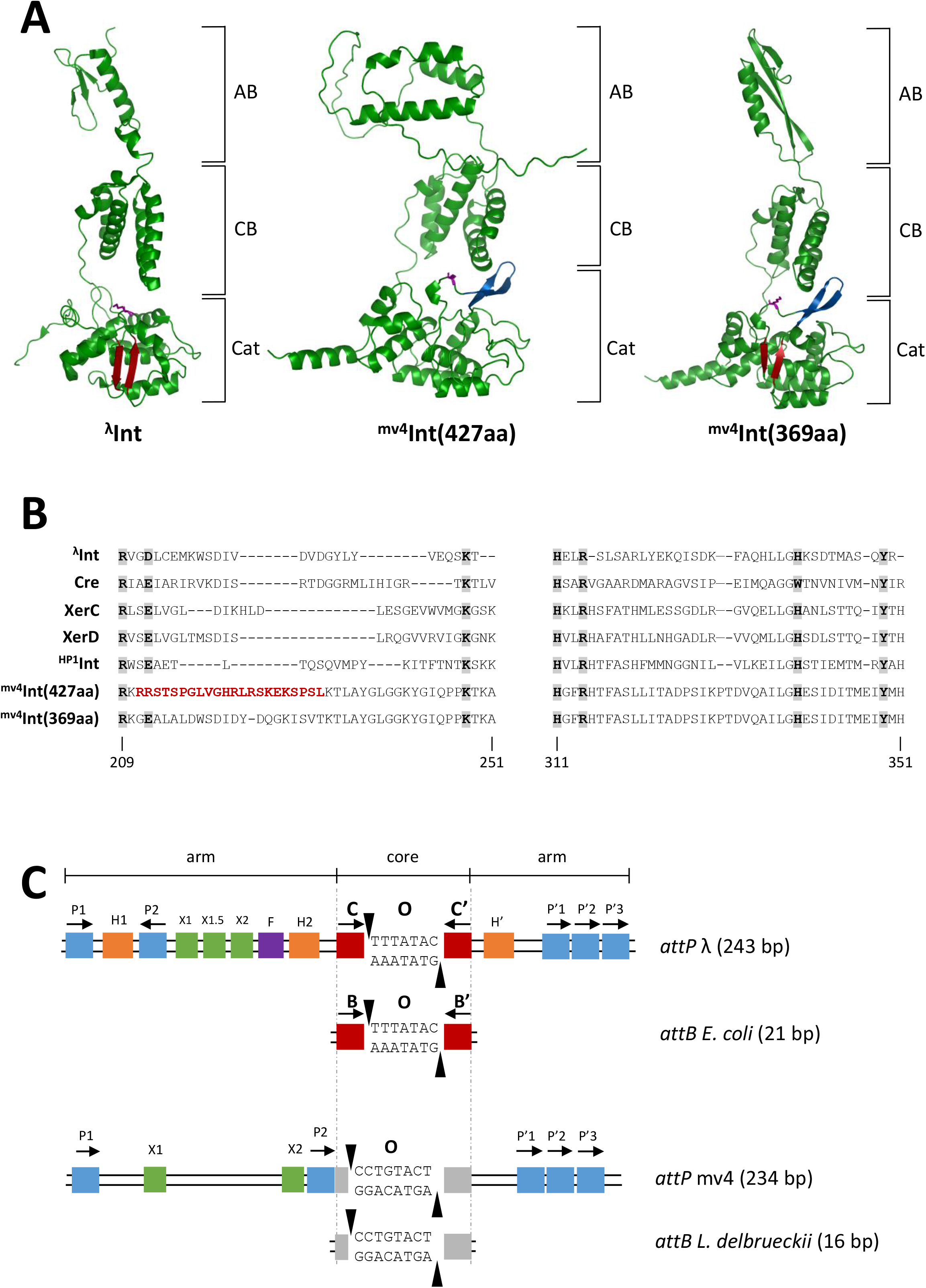
Redefinition of the ^mv4^Int sequence. (**A**) ^λ^Int and ^mv4^Int structure comparison. The ^λ^Int structure (PDB 1Z1G) was taken from Biswas *et al*., (105), and ^mv4^Int(427aa) or ^mv4^Int(369aa) structures were predicted by AlphaFold2 (54). The canonical antiparallel β2- and β3-stands surrounding the catalytic K residue (purple) are indicated in red. The β-hairpin inserted between β2- and β3-strands is highlighted in blue. (**B**) Alignment of catalytic domains of several tyrosine recombinases ^λ^Int, Cre, XerC, XerD, ^HP1^Int, ^mv4^Int(427aa) and ^mv4^Int(369aa). Numbers correspond to aminoacid sequence of ^mv4^Int(369aa). The K residue was manually adjusted according to Nunes-Düby *et al*. alignment (103). The erroneous aminoacids of ^mv4^Int(427aa) are shown in red, and the catalytic domain’s seven conserved residues are highlighted in grey. (**C**) Comparative genetic organisation of lambda (adapted from (13)) and mv4 (adapted from (42)) *attP* and *attB* sites. Arm-binding and core-binding sites for the integrase are represented by blue and red rectangles, respectively. IHF- and Fis-binding sites are represented by orange and purple rectangles, respectively. Excisionase binding sites are shown by green rectangles. Identical sequences between mv4 *attP* and *attB* is shown by grey rectangles. Horizontal black arrows indicate the binding-sites orientation. Vertical black arrows indicate cleavage sites.

Although hundreds of heterobivalent YR recombination systems have been identified in bacterial genomes (25), with each integrase associated with its specific *attP* and *attB* sites, the organisation of their recombination sites has only been fully characterised for a very limited number of systems (Supplementary Figure S2) from *Gammaproteobacteria*, such as phages HP1 from *Haemophilus influenzae* (26), KplE1 and P2 from *E. coli* (27, 28), and P22 from *Salmonella enterica* sv Typhimurium (29). YR recombination systems from only one phage and two ICEs outside *Gammaproteobacteria* has been fully characterised to date: the mycophage L5 (30), Tn916 from *Enterococcus faecalis* (31) and CTnDOT from *Bacteroidetes thetaiotaomicron* (32). Similar to the lambda phage, all recombination reactions rely on intasome formation, and each system contains a ∼ 250-bp complex *attP* site harbouring various binding sites for RDF and the host factors, with the integrase arm- and core-binding sites positioned between them. However, the number of binding sites, their spacing, and orientation generally differ (33), and the intasome structure appears to be characteristic of each Int/*attP* pair (26, 34, 35), underscoring the importance of characterising additional Int/*attP*/*attB* recombination modules to assess the biological diversity of these heterobivalent YR systems.

The temperate bacteriophage mv4 (36, 37) integrates its DNA at the 3’-end of the tRNA^Ser^(CGA) locus of the *Lactobacillus delbrueckii* subsp. *bulgaricus* (hereafter abbreviated as *L. bulgaricus*) chromosome through site-specific recombination (38). Several studies have shown that its recombination module differs from the classical heterobivalent YR systems. It consists of i) a 427-aminoacids heterobivalent integrase (38); ii) a 234-bp *attP* site (39) with a 17-bp region, the core-*attP*, identical to the tRNA^SER^ encoded by the bacteria (Dupont *et al.*, 1995); and iii) a 16-bp *attB* minimal site, the shortest bacterial *attB* site described in the literature (40). The “^mv4^Int/*attP/attB*” system is able to drive recombination in a wide range of bacteria, including *E. coli* and various Gram-positive species (41), indicating that it does not rely on species-specific host-factor to promote recombination. Furthermore, both *attP* and *attB* sites have atypical structures (Figure 1C), with unusual location of ^mv4^Int arm-binding sites on the *attP* site (39, 42), no inverted-repeats core-binding sites at the 17-bp core-*attP* site, and a non-canonical 8-bp overlap sequence (42). As these differences strongly suggested alternative recombination mechanism, we reanalysed the organisation of both *attB* and *attP* sites, through *in vitro* recombination with various random DNA libraries and DNA-binding experiments. We showed that ^mv4^Int is a 369-aminoacids tyrosine recombinase that exhibits structural hallmarks of heterobivalent integrases from the Tn*916* family. We demonstrated that ^mv4^Int is effective on numerous recombination sites as long as they do not deviate for more than one nucleotide from a consensus sequence. We also redefined the organisation of the two recombination sites, demonstrating that *attB* and core-*attP* are longer than expected, contain a 7-bp strand-exchange region, and that their consensus sequences exhibit an organisation similar to the canonical heterobivalent YR recombination sites. Finally, we characterised the different ^mv4^Int binding sites, identifying the *attB* and core-*attP* core-binding sites and refining the locations and length of the arm-binding of *attP*.

## MATERIAL AND METHODS

### Strains, plasmids, oligonucleotides and media

Strains, plasmids and oligonucleotides used in this study are listed in Supplementary Tables S1, S2, and S3. *E. coli* strain MET961 was constructed by replacing the *glgB* gene of strain NEB 5-alpha (New England Biolabs) with the *glgB*::Kan-pWV01*repA* region from *E. coli* EC1000 (43) using the Datsenko and Wanner method (44). *E. coli* strains were grown at 37°C in Lysogenic Broth (45). Antibiotics were used at the following concentration: carbenicillin, 100 µg.ml^-1^; chloramphenicol, 12,5 µg.ml^-1^; erythromycin, 150 µg.ml^-1^; kanamycin, 50 µg.ml^-1^.

### DNA procedures

Standard techniques were used for DNA manipulation and cloning. Polymerase chain reaction (PCR) was performed with Q5-HF polymerase (New England Biolabs, USA) or with CloneAmp Hifi polymerase (Takara Bio, Japan), according to the manufacturer’s instructions. PCR products were purified using the QIAquick PCR purification kit (Qiagen, Germany). Plasmids were constructed either using Gibson assembly (47) with NEBuilder HIFI DNA Assembly (New England Biolabs, USA) or blunt-end cloning with T4 PNK (New England Biolabs, USA) and T4 DNA ligase (New England Biolabs, USA), according to the manufacturer’s instructions. Plasmid DNA was extracted using QIAprep Spin Miniprep kit (Qiagen, Germany) or Nucleobond Xtra Midi (Macherey-Nagel, Germany) and each construction was verified by Sanger sequencing (Mix2seq, Eurofins Genomics, Germany). The different random DNA libraries were constructed as follow: randomized oligonucleotides (109-bp in length for *attB* libraries; 184-bp in length for core-*attP* libraries, Supplementary Table S3B) were obtained by chemical synthesis (Integrated DNA Technologies, USA). PCR was used to create double-stranded DNA using primers attBLib-F / attBLib-R pair for *attB* and attPLib-F / attPLib-R pair for *attP* (Supplementary Table S3). Each PCR-product population was separately cloned by Gibson assembly, either into pCC1FOS™ plasmid (Epicentre, USA) for *attB* libraries, or plasmid pMET359 (Supplementary Table S2) for *attP* libraries. Clones were propagated in *E. coli* strain EPI300 (Epicentre, USA) under chloramphenicol selection for *attB* libraries, or in *E. coli* strain MET961 (Supplementary Table S1) under carbenicillin selection for *attP* libraries.

### Protein Purification

For ^mv4^Int purification, the *E. coli* BL21(DE3) strain containing pMET332 plasmid, (Supplementary Table S2A) was grown in LB at 42°C up to an OD_600_ of 0.6. Integrase gene expression was induced by addition of 0.1 mM of IPTG, and the culture was incubated at 22°C for 3h. Cells were recovered by centrifugation, resuspended in buffer A (50 mM Tris pH 8, 500 mM NaCl, 20 mM imidazole, 10% glycerol) supplemented with 1 mg/ml lysozyme and one tablet of SIGMAFAST Protease Inhibitor Cocktail Tablets EDTA-Free (Merck KGaA, Germany), and disrupted by sonication (10 cycles of 30 sec at 40 % intensity in ice, with 45 sec pause between each cycle). The lysate was cleared by centrifugation (20.000 g, 4°C, 20 min). ^mv4^Int was purified on nickel-nitrilotriacetic acid affinity resin (1ml His-trap HP, GE Healthcare, USA). Column equilibration was performed by injecting 10 column volumes of buffer A. After equilibration, the lysate was injected and unbound proteins were washed using 10 column volumes of buffer A. ^mv4^Int was eluted using a gradient from 0 to 30 % of buffer B (50 mM Tris pH 8, 500 mM NaCl, 500 mM imidazole, 10 % glycerol). Eluted fractions were then injected in a gel filtration column (HiLoad 16/60 Superdex 200, GE Healthcare, USA). This column was equilibrated using 2 column volumes of buffer C (50 mM Tris pH 8, 500 mM NaCl, 10 % glycerol, 1 mM DTT, 1 mM EDTA) and the fractions containing ^mv4^Int were injected and eluted using the same buffer. Eluted fractions containing ^mv4^Int were then 2-fold diluted in buffer D (50 mM Tris pH 8, 10 % glycerol, 1 mM DTT, 1 mM EDTA). Fractions containing ^mv4^Int were injected on heparin column (1ml HiTrap Heparin HP, GE Healthcare, USA) equilibrated with 10 column volumes of buffer E (50 mM Tris pH 8, 250 mM NaCl, 20% glycerol, 1 mM DTT, 1 mM EDTA), and unbound proteins were removed using 10 column volumes of buffer E. ^mv4^Int was eluted using a gradient from 0 to 100% of buffer F (50 mM Tris pH 8, 1 M NaCl, 20 % glycerol, 1 mM DTT, 1 mM EDTA). The different purification steps were monitored by SDS-PAGE (Supplementary Figure S3A). Purified integrase was aliquoted, snap-frozen in liquid N2 and stored at -80°C in buffer containing 50 mM Tris pH 8, 500 mM NaCl, 20% glycerol, 1 mM DTT, 1 mM EDTA.

For 6xHis-HU_Lla_ purification, the *E. coli* BL21(DE3) strain containing pMET369 plasmid (Supplementary Table S2) was grown in LB at 37°C up to an OD_600_ of 0.6. Expression of the *hup* gene was induced by addition of 0.4 mM of IPTG and the culture was incubated at 22°C for 3h. Cells were recovered by centrifugation, resuspended in buffer A, and disrupted by sonication as indicated above. The lysate was cleared by centrifugation (16.000 g, 4°C, 15 min), and the recovered supernatant was heated at 70°C for 30 min and cleared by centrifugation (16.000 g, 4°C, 15 min). The 6xHis-HU_Lla_ protein was purified on nickel-nitrilotriacetic acid affinity resin as described for ^mv4^Int, except than elution was performed with a gradient from 0 to 70% of buffer B. Buffer of the eluted fractions containing 6xHis-HU_Lla_ was replaced by buffer G (50 mM Tris pH 8, 200 mM NaCl) in Amicon® Ultra – 3 kD Centrifugal Filters (Merck KGaA, Germany) and resulting solution was injected in a heparin column. Unbound proteins were removed using 10 column volumes of buffer E, and 6xHis-HU_Lla_ protein was eluted using a gradient from 0 to 100% of buffer F. The different purification steps were monitored by SDS-PAGE (Supplementary Figure S3A). Purified HU was aliquoted, snap-frozen in liquid N2 and stored at -80°C in buffer containing 50 mM Tris pH 8, 150 mM NaCl.

### *In vitro* fluorescent assay

Reaction mixtures (20 µl) contained 0.08 pmol (300 ng) of supercoiled plasmid carrying the *attP* site, 0.08 pmol (15 ng) of linear fluorescent (Cy3), ∼300-bp *attB* fragment obtained by PCR amplification with LbbulgattB-F / Cy3-New-attB-R primers pair, 7.2 pmol (300 ng) of ^mv4^Int, and 40 µg of a crude-extract from *E. coli* BL21(DE3) heated at 95°C for 10 min, in TENDP 1X buffer (25 mM Tris pH 7.5, 1 mM EDTA, 150 mM NaCl, 1 mM DTT, and 10% PEG8000). The reaction was incubated at 42°C either 1h30 or 16h, depending on the presence or absence of the heated *E. coli* crude-extract respectively, and was stopped by addition of 0.1% SDS. Samples were analysed by electrophoresis in 0.8% agarose gels. Fluorescence was revealed using the ChemidocMP imaging system (Bio-Rad, USA).

### *In vitro* recombination assay of random DNA libraries

The reaction (20 µL) was performed with 0.08 pmol of *attB*LibX plasmid (X being a number from 5 to 9, Supplementary Table S3) or *attP*LibX plasmid (X being a number from 1 to 4, Supplementary Table S3), 0.08 pmol of pBS*attB* or pMC1 plasmid (38), 7.2 pmol of ^mv4^Int and 40 µg of crude-extract from *E. coli* BL21(DE3) heated at 95°C for 10 min, in TENDP 1X and incubated 1h30 at 42°C. The different regions *attB*, *attP*, *attL* and *attR* were amplified by PCR using SeqLibattB-F / SeqLibattB-R primers pair for *attB*; SeqLibattP-F / SeqLibattP-R primers for *attP*; SeqLibattB-F / SeqLibattL-R primers for *attL* and SeqLibattR-F / seqLibattP-R primers for *attR*. PCR products were purified and either analysed by Sanger sequencing (Mix2seq, Eurofins Genomics, Germany) or Next-Generation Sequencing (Illumina, NGSelect Amplicon, 2 × 150 bp, Eurofins Genomics, Germany).

### Bioinformatic analyses

For NGS analyses, reads from the two fastq files were paired using CLC Genomics Workbench 7 (Qiagen, Germany). Paired and unpaired data were uploaded and analysed on the public server at usegalaxy.org (48). First, the samples were sorted by length using “Filter Fasta” program (Galaxy version 2.3). Then, unwanted sequences in each group (*attB*, *attP*, *attL* and *attR*) were removed with “Bowtie2” program (49), using the expected sequence for each group as the reference genome. For each group, “Fasta Width formatter” program (Galaxy version 1.0.1) was used to get each fasta sequence on a single line. Sequences containing ambiguous bases were removed by using “seqtk_seq” program (Galaxy version 1.3.3). For each group, forward and reverse sequences were separated in two different files using “Filter Fasta” program. Reverse sequences were reverse-complemented using “seqtk_seq” program, and merged with the forward sequences using “merge.files” program (50). To eliminate any contaminant sequence, a final verification was performed in each group using “Filter Fasta” program, by keeping only reads that perfectly matched the expected sequence, and heptamers corresponding to the randomized region of DNA libraries were extracted from each sequence using the “Trim sequences” program (Galaxy version 1.0.2), and their occurrence calculated using the “Wordcount” program (51). Only heptamers present in the two biological replicates were considered for further statistical analyses, such as mutual information (52) and Cramer’s V (53) analyses, with occurrence values corresponding to the mean of the two replicates.

### Electrophoretic Mobility Shift Assay

HPLC-purified 5’ Cy3 end-labelled synthetic oligonucleotides were obtained by chemical synthesis (Eurofins Genomics, Germany). Labelled dsDNA substrates were prepared by hybridization of 200 pmol of labelled and 300 pmol of unlabelled complementary oligonucleotides (Supplementary Table S3C) in 10 mM Tris pH 7.5, 50 mM NaCl, and incubating the samples at 95°C for 5 min in a thermal cycler (Bio-Rad, USA) and decreasing the temperature of 1.5°C/min until it reaches 25°C. Binding reactions (20 µl) were performed with 0.87 pmol of labelled dsDNA and 4.48 pmol of unlabelled dsDNA in buffer containing 25 mM Tris pH 8, 75 mM NaCl, 10% glycerol, 0.5 mM DTT, 0.5 mM EDTA, 1 µg polydIdC (Merck KGaA, Germany), 0.1 mg/ml BSA. The protein was added, the reaction performed at room temperature for 20 min, and samples were loaded onto a non-denaturing 7.5% polyacrylamide gel (Mini-PROTEAN TGX, Bio-Rad, USA). Gels were run at 4°C, 75V for 2 h. Fluorescence was revealed using the ChemidocMP imaging system (Bio-Rad, USA).

## RESULTS

### The ^mv4^Int is a 369-aminoacids tyrosine integrase

The previous analyses of mv4 phage’s integration module described the ^mv4^Int as a 427-aminoacids (aa) protein with significant similarity with the ^λ^Int integrase (38), although ^mv4^Int contained only six of the seven conserved residues of YRs catalytic pocket, with the structurally important D/E residue missing (Figure 1B). However, when ^mv4^Int(427aa) was subjected to protein structure prediction program (54), discrepancies were observed when compared to classical heterobivalent integrases (Figure 1A), such as the absence of the encompassing β2 and β3 strands around the catalytic pocket’s lysine residue (K235 for ^λ^Int, Aihara *et al.*, 2003) as well as the deficiency of a structured arm-binding domain (56). The *int* gene from the pMC1 plasmid (38) was then resequenced, revealing five single nucleotide deletions and two inversions (Supplementary Figure S1A) compared to the initially published sequence. These nucleotides modifications were further validated in other *int* gene-containing plasmids, such as pA3int and pET-Int (39), as well as from the PCR amplicon of the *int* region of mv4 prophage (data not shown). These nucleotide changes exerted a profound impact on the ^mv4^Int aminoacid sequence, such as truncation of the protein to 369 aminoacids and divergence by 53 aminoacids from the published integrase. This revised protein profile aligns identically or very closely with prophage integrases identified within the genomes of multiple *L. bulgaricus* strains (Supplementary Figure S1B), and comparison of its catalytic domain (residues 172 to 369) against ^λ^Int, Cre, XerC, XerD or ^HP1^Int recombinases indicates the inclusion of all the conserved residues from YRs catalytic pocket (Figure 1B). In addition, its predicted structure now displays a well-structured arm-binding domain, as well as the canonical antiparallel β2-β3-strands (red arrows, Figure 1A). Interestingly, ^mv4^Int contains two additional antiparallel β-strands (blue arrows, Figure 1A) positioned between the β2-strand and the catalytic lysine, a structural hallmark from the ^Tn*916*^Int subgroup (25) required for DNA cleavage and strand-exchange reaction as demonstrated for ^Tn*1549*^Int (57). The ^mv4^Int(369aa) protein was overproduced, purified (Supplementary Figure S3A), and used for *in vitro* recombination assays between the mv4 *attP* site located on a supercoiled plasmid (pMC1, Supplementary Table S2) and a fluorescent 290-bp PCR product derived from the *L. bulgaricus* tRNA^SER^(CGA) locus (38). During the experimental optimisation of this *in vitro* assay with wild-type (WT) or modified *attP*/*attB* sites, a decrease in recombination activity was observed if using purified ^mv4^Int instead of *E. coli* cell extract overexpressing the integrase, particularly for the modified sites (Supplementary Figure S3B, middle *vs* left gel) However, addition of crude extract from *E. coli* or *Lactococcus lactis* (data not shown) led to a notable enhancement in recombination activity, with improvements by factors of 2-3 for WT sites and 25-30 for modified sites (Supplementary Figure S3B, right). Both crude extracts retained their efficacy even after heat treatment but not after proteinase K treatment, indicating that the “stimulating factor” could be attributed to a heat-resistant protein present in *E. coli* and *L. lactis* cells. As the only conserved Nucleoid -Associated Protein (NAP) between the two species is the HU protein, an ubiquitous protein found in all prokaryotic phyla (58), we overproduced and purified HU protein from *L. lactis* (Supplementary Figure S3A). Addition of purified HU in the reaction superseded the stimulation observed with the crude extract in a dose dependent manner (Supplementary Figure S3C), demonstrating that HU is not *stricto sensu* a host factor for ^mv4^Int-mediated recombination, such as IHF or Fis for ^λ^Int, but acts as a facilitator for recombination, particularly in the context of modified recombination sites. Finally, we unequivocally demonstrated that the purified ^mv4^Int(369aa) catalyses the site-specific recombination without *L. bulgaricus* host factor (Supplementary Figure S3D), and showed that HU protein acts as a facilitator for recombination, particularly in the context of modified recombination sites. We also demonstrated that recombination was abolished, or below the detection limit of the method, when using variants of two important residues (Y349F or K248A, Supplementary Figure S3D) of the YR catalytic pocket.

### Global strategy of the use of random DNA libraries

Given the organisational peculiarities of the mv4 *attP* core-region and *attB* site (40, 42), both sites underwent a comprehensive experimental reevaluation using a random DNA libraries based approach (59), with the rational that DNA positions encompassing these core regions should display constraints in nucleotide composition. Those DNA libraries, corresponding either to the *attB* site or to the core-region of the *attP* site, harboured a range of 7 to 10 randomized positions (Figure 2A). For that purpose, oligonucleotides containing the four possible nucleotides at determined positions were synthetized, amplified by PCR and cloned in *E. coli* into different plasmids (pMET359 for core-*attP* libraries, and pCC1FOS for *attB* libraries, Supplementary Table S2). Each plasmid library was recovered and used for *in vitro* recombination experiments with either the native partner site (*attP*_WT_ or *attB*_WT_) or the associated partner library (i.e. two libraries with the same randomized region for the 2 sites). After recombination, the *attL* and *attR* sites were amplified by PCR and sequenced. Among the randomized positions, only nucleotides permissive for recombination are expected to be retrieved in these hybrid sites, which make it possible to determine the constraints exerted on the nucleotide composition at each evaluated position (Figure 2B).

**Figure 2.**
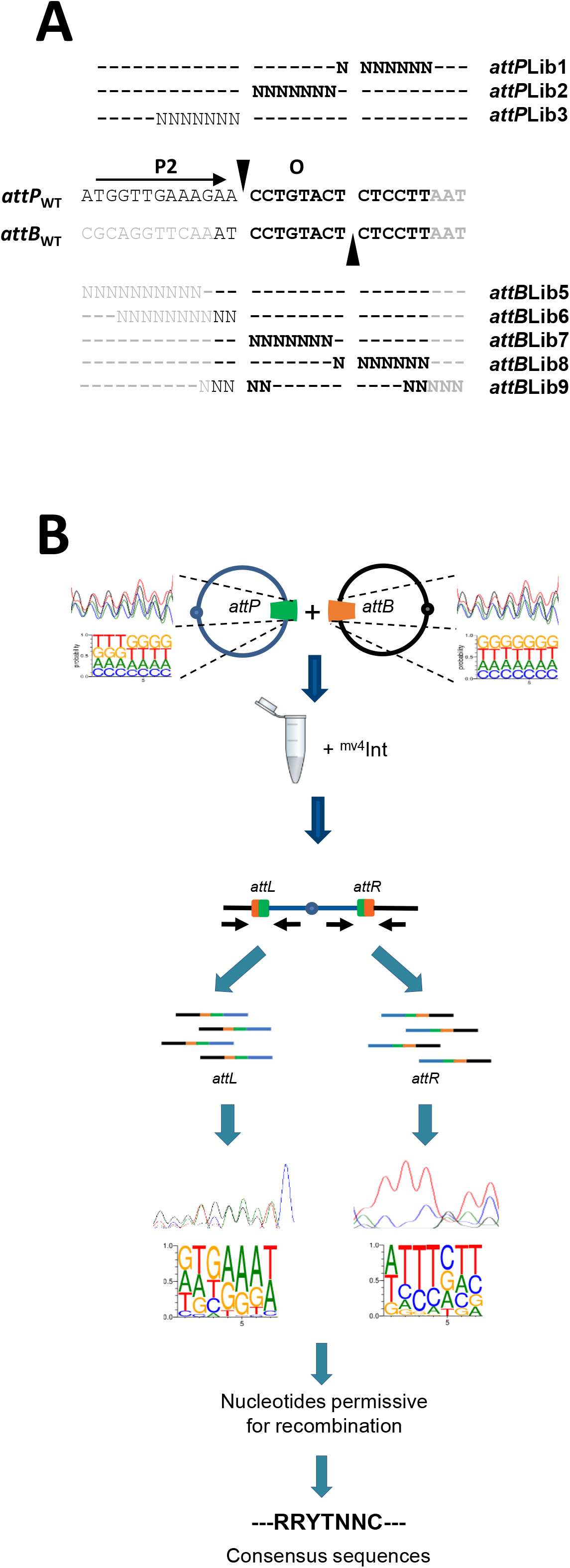
Use of randomized libraries for the characterization of core-*attP* and *attB sites*. (**A**) Localization of random nucleotides in core-*attP* or *attB* sites. Nucleotides outside the *attB* minimal site (40) are indicated in grey, and sequence identical between core-*attP* and *attB* are in bold. The atypical P2 arm-type binding site and strand-exchange (O) region (42) are indicated by horizontal and vertical black arrows, respectively. Random nucleotides are represented by “N” and nucleotides identical to the native sequences are symbolized by dashes. (**B**) Global strategy for the use and analyses of randomized libraries (see text). The size of the letters in the *Sequence Logos* indicates the proportion of each nucleotide at each position. The consensus sequence indicated is arbitrary.

### The mv4 *attB* is a 21-bp site

The *attB* minimal site has previously been characterized as a 16-bp site (Figure 3A, Auvray *et al.*, 1999b). However, when a DNA fragment containing this 16-bp sequence surrounded by nucleotides different from the *L. bulgaricus* chromosome (Figure 3A) was tested by *in vitro* recombination against a plasmid-borne *attP*_WT_ site, no recombination could be observed (Figure 3B), implying that the 16-bp fragment did not contained the complete native *attB* site. In contrast, a recombination product was recovered with a 23-bp fragment corresponding to the minimal *attB* with additional native *L. bulgaricus* nucleotides at its left (Figure 3A and 3B). In order to delineate the boundaries of the *attB* site, a DNA library, made of five randomized positions overlapping each end of the published *attB* site, was constructed (*attB*Lib9, Figure 2A) and subjected to *in vitro* recombination against the *attP*_WT_ site. Upon recombination, the resultant *attL* and *attR* sites exhibited different degrees of nucleotide degeneracy indicative of the constraints exerted at the left and the right sides of the *attB* site, respectively (Figure 3C). Among the five positions at the right side of *attB*, only the first two positions exhibited constraint in their nucleotide composition (no G in pos6 and no A in pos7, Figure 3C), thereby confirming that the *attB* site ends with the sequence 5’-CTCCTT-3’, as previously published (40). In contrast, all five positions at the left side of the *attB* site (pos1 to pos5, Figure 3C) displayed strong constraints, even those previously identified outside of the minimal *attB* site (40). Subsequently, the left boundary of the *attB* site was confirmed by recombination between the *attP*_WT_ site and two additional *attB* random libraries (*attB*Lib5 and *attB*Lib6, Figure 2A), each containing ten randomized nucleotides overlapping the left end of the permissive 23-bp fragment. Both cases yielded a lack of constraints up to the second G of the sequence (pos4 Figure 3D, and pos7 Figure 3E), thereby implying that the mv4 *attB* site corresponds to a 21-bp site of sequence 5’-TTCAAATCCTGTACTCTCCTT-3’.

**Figure 3.**
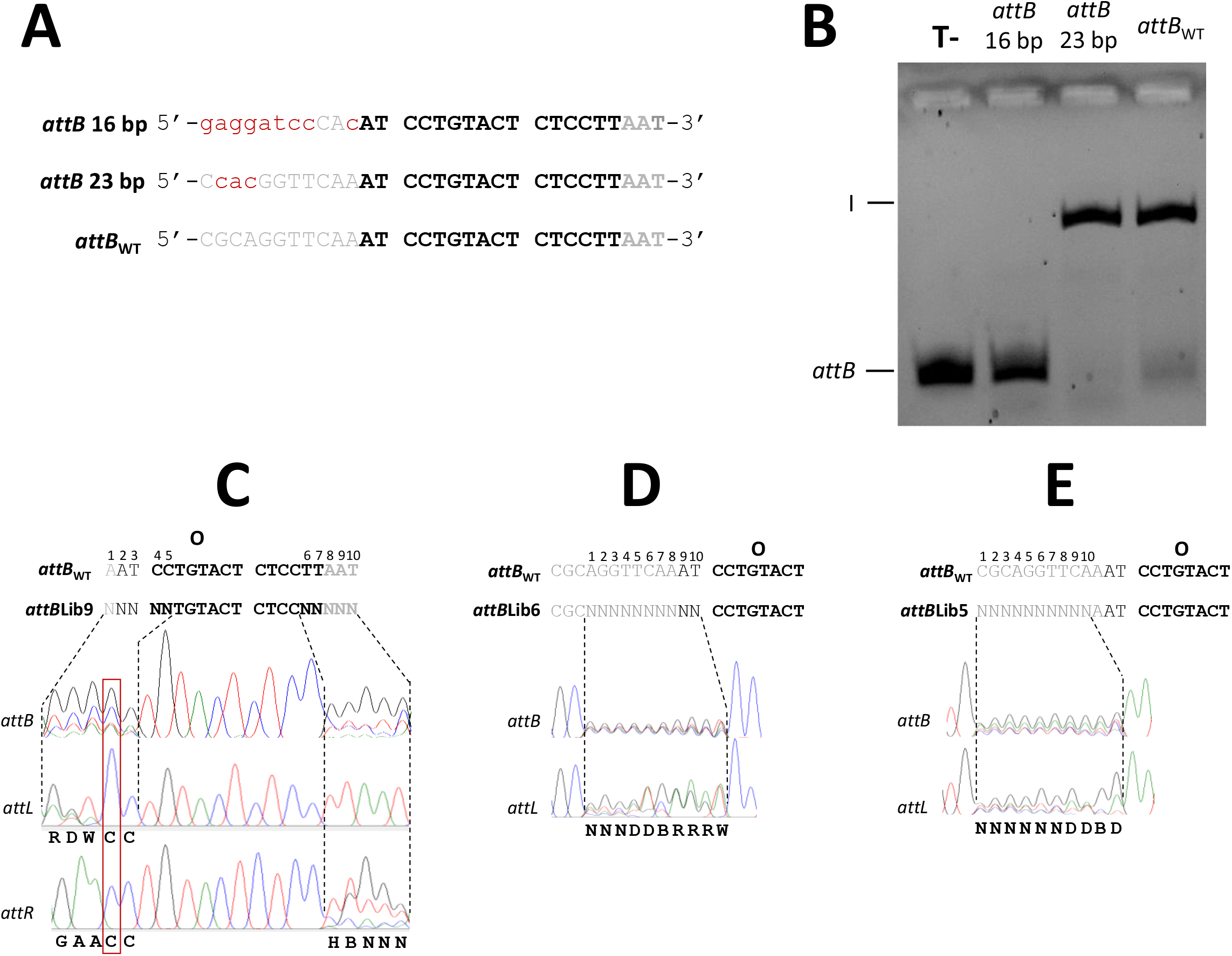
*In vitro* characterization of the minimal *attB* size. (**A**) DNA sequences used for the *attP*_WT_/*attB in vitro* recombination assay. *attB* site is described as in Figure 2A. Nucleotides differing from the *L. delbrueckii* subsp *bulgaricus* sequence are indicated in red lowercase letters. (**B**) *attB* size-dependent effect on the recombination reaction. The size is indicated above each lane. The fluorescent *attB* fragment and the linear recombination product (I) are indicated as in Supplementary Figure S3. T-, control (reaction without ^mv4^Int). (**C**) Chromatogram of *attB*, *attL* and *attR* regions from *attB*Lib9 × *attP*_WT_ *in vitro* recombination (n=2). IUPAC code, R= A/G, D=A/G/T, W=A/T, H=A/C/T, B=C/G/T. (**D**) Chromatogram of *attB*, and *attL* regions from *attB*Lib6 × *attP*_WT_ recombination (n=3). (**E**) Chromatogram of *attB*, and *attL* regions from *attB*Lib5 × *attP*_WT_ recombination (n=2).

### The recombination specificity depends on a 7-bp overlap region

A previous study based on the use of DNA suicide substrates determined the length of the *attB*_mv4_ and *attP*_mv4_ overlap regions to 8 bp (42), i.e. within the size-range characteristic of YR recombination systems (5). However, we noticed that heterobivalent recombination systems characterized to date (29, 60–64) generally exhibit a 7-bp overlap region. In addition, it has been soon observed that nucleotide composition in the *attP*/*B* overlap regions was not important for recombination, but that sequence identity between the two sites was mandatory (65–69), though heteroduplexes can sometimes be functional (59, 70, 71). Leveraging these properties, we re-evaluated the size and constraints governing the overlap region of the mv4 recombination system by performing *in vitro* recombination with different random *attP* or *attB* libraries. Size of the overlap region was determined with the rational that if randomized position of the library resides within the overlap region, only nucleotide identical to the *attP*_WT_ or *attB*_WT_ site would be recovered in both *attL* and *attR* sites (Figure 4A). Conversely, if the randomized position is located outside the overlap region, *attL* and *attR* sites would exhibit different nucleotides; one site with the wild-type nucleotide and the other with the permissive nucleotides from the library (Figure 4A). The previously determined left border (42) was validated through a*ttB*Lib9 × *attP*_WT_ recombination, which yielded only the WT sequence at position 4 and 5 of both *attL* and *attR* sites (red box, Figure 3C). In contrast, examination of the right border through *attP*Lib1 × *attB*_WT_ or *attB*Lib8 × *attP*_WT_ recombinations yielded different *attL* and *attR* sites for each case (Figure 4B), indicating that the position 1 of these random libraries is unlikely to localize within the overlap region. It was thereby determined that the core-*attP* and *attB* overlap regions are 7-bp in length, consistent with the characteristic size observed for other heterobivalent YR systems.

**Figure 4.**
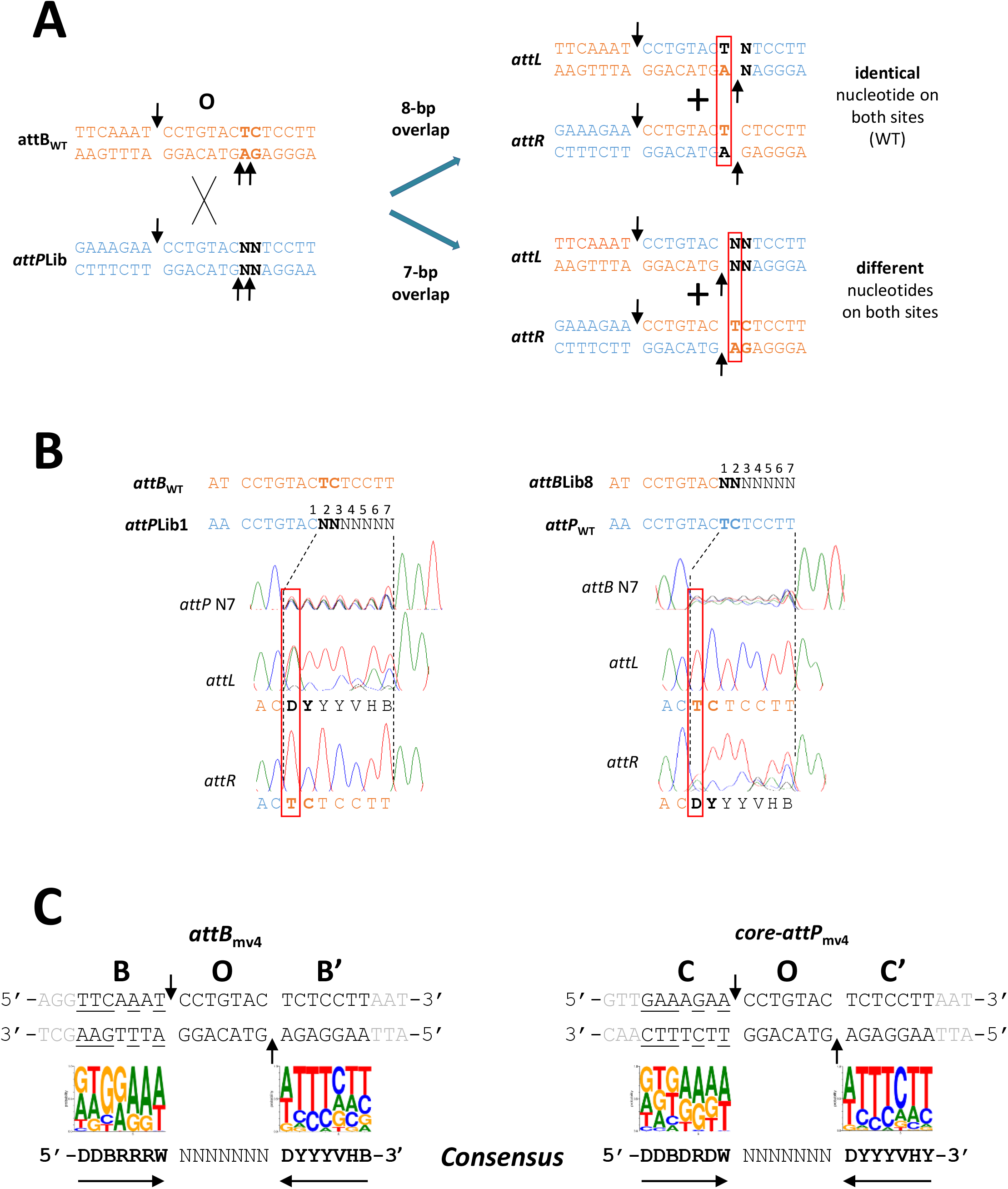
Characterisation of the strand-exchange (overlap) region. (**A**) Expected sequence of recombination products depending on cleavage site position. For an 8-bp overlap region, both *attL* and *attR* should contain the WT nucleotide (T) at the first position (boxed in red) of the randomized region. For a 7-bp region, sequence at the first position should be different between *attL* and *attR*, with all permissive nucleotides present at one of the recombined sites. (**B**) Chromatograms of *attP, attB*, *attL,* and *attR* regions from *attB*_WT_ × *attP*Lib1 recombination (n=3), or *attB*, *attL, attR* regions from *attP*_WT_ × *attB*Lib8 recombination (n=3). (**C**) *Sequence Logos* from NGS sequencing revealing biased nucleotide of recombination products composition in regions surrounding the overlap. Only sequencing reads present at least 10 times with a tenfold enrichment were considered for the analysis. Nucleotides outside *attB* or core-*attP* sites are indicated in grey, and those different between *attB* and core-*attP* are underlined.

Recombination specificity was determined through *in vitro* recombinations between one WT site (*attB* or *attP*) and cognate DNA libraries containing random nucleotides (*attB*Lib7 × *attP*_WT_, or *attP*Lib2 × *attB*_WT_, Figure 2A). Following recombination and subsequent sequencing of *attL* and *attR* sites, only the WT overlap sequence was recovered among the 16,384 (4^7^) theoretical combinations (Supplementary Figure S4A and S4B). This underscores the strict requirement for identical overlap regions for a complete recombination reaction. Recombination assays between the two random libraries (*attB*Lib7 × *attP*Lib2) revealed the recovery of all possible nucleotides at each randomized position (Supplementary Figure S4C). Finally, recombination of *attB*Lib7 against three *attP* sites with distinct overlap sequence, yielded only the sequence identical to the *attP* tested (Supplementary Figure S4D), though recombination on sequences containing heteroduplexes could not be totally excluded (Supplementary Figure S4D, left). However, as already observed for model YRs such as ^λ^Int (72), Flp (73, 74), or Cre (59), the composition of the overlap region can significantly influence the recombination efficiency, since *in vitro* fluorescent assays with *attB* sites adapted to each *attP* overlap region (Supplementary Figure S4E) revealed the formation of recombined products from level comparable to WT sites (*attP*_1_/*attB*_1_ and *attP*_2_/*attB*_2_ pairs) to nearly the detection limit of the method (*attP*_3_/*attB*_3_). Altogether, these results suggest a limited set of constraints on the nucleotide composition of the mv4 *attB*/*attP* overlap regions, implying that ^mv4^Int can catalyse recombination across any nucleotide combination, with variable efficiencies, as long as the core-*attP* and *attB* overlap regions are identical.

### *attB* and core-*attP* organisations are similar to classical YR recombination sites when considering nucleotide degeneracy

The *attB*Lib6 library, which originally contributed to the determination of the *attB* site’s size, also unveiled notable nucleotide constraints on the left side of the *attB* overlap region, since after recombination against *attP*_WT_, some nucleotides were absent at seven positions (from pos 4 to 10, Figure 3D). Similar constraints were observed at corresponding positions when analysing products resulting from *attP*_WT_ × *attB*Lib9 and *attP*_WT_ × *attB*Lib5 recombinations (Figure 3C and 3E, respectively), demonstrating absence of experimental bias when moving the randomized region’s windows across the recombination site. These results imply that *attB* site can support nucleotide variations on its left-side when recombined with *attP*_WT_, thereby suggesting the possibility to define the degenerated sequence 5’-DDBRRRW-3’ as being permissive for *in vitro* recombination. Similarly, the exploration of the right side of the core-*attP* and *attB* sites (Figure 4B and 4C) led to the definition of the degenerate sequence 5’-DYYYVHB-3’ as permissive for recombination. This indicates that ^mv4^Int’s interaction with its recombination sites is adaptable enough to tolerate some nucleotide variation. To better characterize this flexibility, DNA libraries containing 7-bp random nucleotides on each side of the *attB* and core-*attP* overlap regions (*attB*Lib6, *attB*Lib8, *attP*Lib3, and *attP*Lib1, Figure 2A) were subjected to Next-Generation Sequencing (NGS), as well as the *attL* or *attR* sites obtained after recombination against *attP* or *attB* wild-type sites, depending on the library. Each library contained between 99.5% (16312) and 100% (16384) of the theoretically expected number of heptamers (4^7^). However, after recombination, 33% to 72 % of the DNA motifs were retrieved on *attL* or *attR* sites (Supplementary Table S4), confirming that only a subset of the recombination sites was permissive for recombination reaction. The *Sequence Logos* generated for each *attB* or *attP* library demonstrated near-uniform nucleotide frequencies at each position, with G and T prevailing over A and C and a slight tendency for cytosines to be underrepresented (Supplementary Figure S5A). However, after recombination, *attL* and *attR* recombined sites clearly exhibited strong biases in nucleotide distribution. The biases were very similar for *attP* and *attB*, with enrichment in G and A at the left-side region and in C and T at the right-side (Supplementary Figure S5A). Furthermore, these *Sequence Logos* confirmed the nucleotide degeneracy observed by Sanger sequencing of *attB* and core-*attP*, with additional nucleotides present at lower frequencies due to the highest resolution of NGS that might contribute to background noise amplification. As it is highly improbable that all motifs from the libraries are functional, we assumed that comparing heptamers counts before and after recombination (i.e. calculating an enrichment ratio) could yield meaningful insights into their potential to form a productive recombination complex with ^mv4^Int. Initial observations revealed highly skewed read counts for each motif after recombination, ranging from 69525 to 1 (Supplementary Table S4). Moreover, the relationship between read counts and their rank exhibited rapid decline (Supplementary Figure S5B), with only 2.8% to 7.5% of heptamers representing 50% of the total read counts (Supplementary Table S4), and 31.5% to 52.4% of heptamers occurring fewer than 10 times (data not shown). Additionally, two independent trials resulted in hundreds of heptamers present in only one of the experiments, leading us to consider as informative only the 4842 (*attB*Lib6 × *attP*_WT_), 8472 (*attB*Lib8 × *attP*_WT_), 9973 (*attB*_WT_ *× attP*Lib3), and 5699 (*attB*_WT_ *× attP*Lib1) heptamers present on both assays (Supplementary Table S4). Finally, we normalized the nucleotide prevalence at each randomized position through mutual information (MI) analyses (52) (Supplementary Figure S5C), and estimated the minimum read count and enrichment ratio appropriate to mitigate background noise from *Sequence Logos* and facilitate subsequent statistical analyses. Read count and enrichment ratio thresholds of 10 allowed to retain nearly half of the information (read count) for each library, except for *attL* site from *attB*_WT_ × *attP*Lib3 recombination where a threshold of 6 in enrichment ratio was necessary. Altogether, this led us to consider for the further analyses “reduced” datasets corresponding to 117 heptamers from *attB*Lib6 × *attP*_WT_ recombination, 613 from *attB*Lib8 × *attP*_WT_ recombination, 1025 from *attB*_WT_ × *attP*Lib3 recombination, and 231 from *attB*_WT_ × *attP*Lib1 recombination. The *Sequence Logos* generated (Figure 4C) corroborated previous consensus sequences and unequivocally demonstrated that ^mv4^Int has the ability to recombine a wild-type *attP* or *attB* site with its counterpart displaying identical overlap region but with some nucleotide variation either on the left- or at its right-side. Interestingly, when considering the consensus sequences rather than the WT sequences, the DNA motifs surrounding the overlap region exhibited complementarity, with purines enrichment at the left-side and pyrimidines at the right-side. Thus, in contrast to the previous study that concluded there might be an arm-type site adjacent to the overlap region (42), our analysis strongly supports the classical organization of heterobivalent YR recombination systems for the 21-bp *attB* and core-*attP* sites, with imperfect inverted-repeated patterns flanking each strand-exchange region (Figure 4C), and defining BOB’ and COC’ structures for *attB* and core-*attP* respectively, in accordance with the phage lambda terminology.

### Identifying the ^mv4^Int core- and arm-binding regions of *attP* and *attB* sites

In the classical heterobivalent YR recombination systems, the sequences surrounding the overlap region define the core-binding sites of the integrase (28, 32, 75). To demonstrate that B, C and B’/C’ regions correspond to the ^mv4^Int core-binding sites, different gel-shift assays were performed. For the model lambda recombination system, it has been demonstrated that the integrase core-binding domain is only able to bind to the core-*attP* (COC’) if its arm-binding domain is bound to the *attP* arm sites (75). We thus first compared the ^mv4^Int binding to a labelled 35-bp dsDNA fragment containing the 21-bp COC’ core-*attP* region (Supplementary Table S3C), in the presence or absence of unlabelled dsDNA containing the P’1P’2 arm sites (Supplementary Table S3C) characterized previously (39). As for lambda, the ^mv4^Int/COC’ complex exhibited instability in absence of the P’1P’2 fragment, resulting in faint bands and a strong background smear (Figure 5A). However, upon adding a 28-bp dsDNA containing the P’1P’2 sequences, ^mv4^Int binding to the core-*attP* sequence was strongly stabilized and led to the formation of three complexes (Figure 5A). The direct effect of P’1P’2 sites on interaction of ^mv4^Int with the core-binding sites was clearly observed by the use of a 40-bp dsDNA containing the P’1P’2 sites, which revealed a shift in mobility of complexes C2* and C3* (Figure 5A), compared to those observed with the 28-bp fragment. Interestingly, ^mv4^Int was also able to stably bind to a labelled P’1P’2 fragment only upon addition of unlabelled COC’ fragment (Supplementary Figure S6A), suggesting that AB and CB domains exert a reciprocal inhibition on their binding at the *attP* site, indicative of a mechanism governing intasome formation distinct from lambda (75) or CTnDOT (32). According to studies from other YR systems, our data suggest that complex C1 likely corresponds to a single ^mv4^Int monomer bound to one core-binding site, while complex C2 is indicative of one monomer bound to both an arm and a core-binding site, and complex C3 corresponds to ^mv4^Int dimer bound to arm- and core-binding sites simultaneously. In presence of unlabelled P’1P’2 fragment, ^mv4^Int stably binds *in vitro* with the same efficiency to both core-*attP* and *attB* sites in a dose-dependent manner (Figure 5B). As the left and right half-sites of *attB* and core-*attP* are not perfect inverted repeats, a feature that can be relied to asymmetric binding of the Y recombinases to their cognate recombination sites (28, 76), gel shift assays were conducted using labelled 35-bp dsDNA fragments containing individual half-sites (B’/C’, B, or C) (Supplementary Table S3C). The binding complexes C1, C2, and C3 observed with *attB*/core-*attP* site were also formed with the B’/C’ half-site arm (Supplementary Figure S6B), albeit at lower efficiency compared to the WT site, especially for the potential dimer bound to both arm- and core-binding sites (C3). Moreover, ^mv4^Int bound to the B or C half-sites with significantly lower affinity than the B’/C’ half-site, as only complex C1 was detectable (Supplementary Figure S6B). Thus, ^mv4^Int appears to efficiently bind to both the core-*attP* and *attB* sites in a cooperative manner, with a preference for the conserved right half-site (B’/C’), most likely promoting the binding of the second ^mv4^Int monomer at the B or C sites.

**Figure 5.**
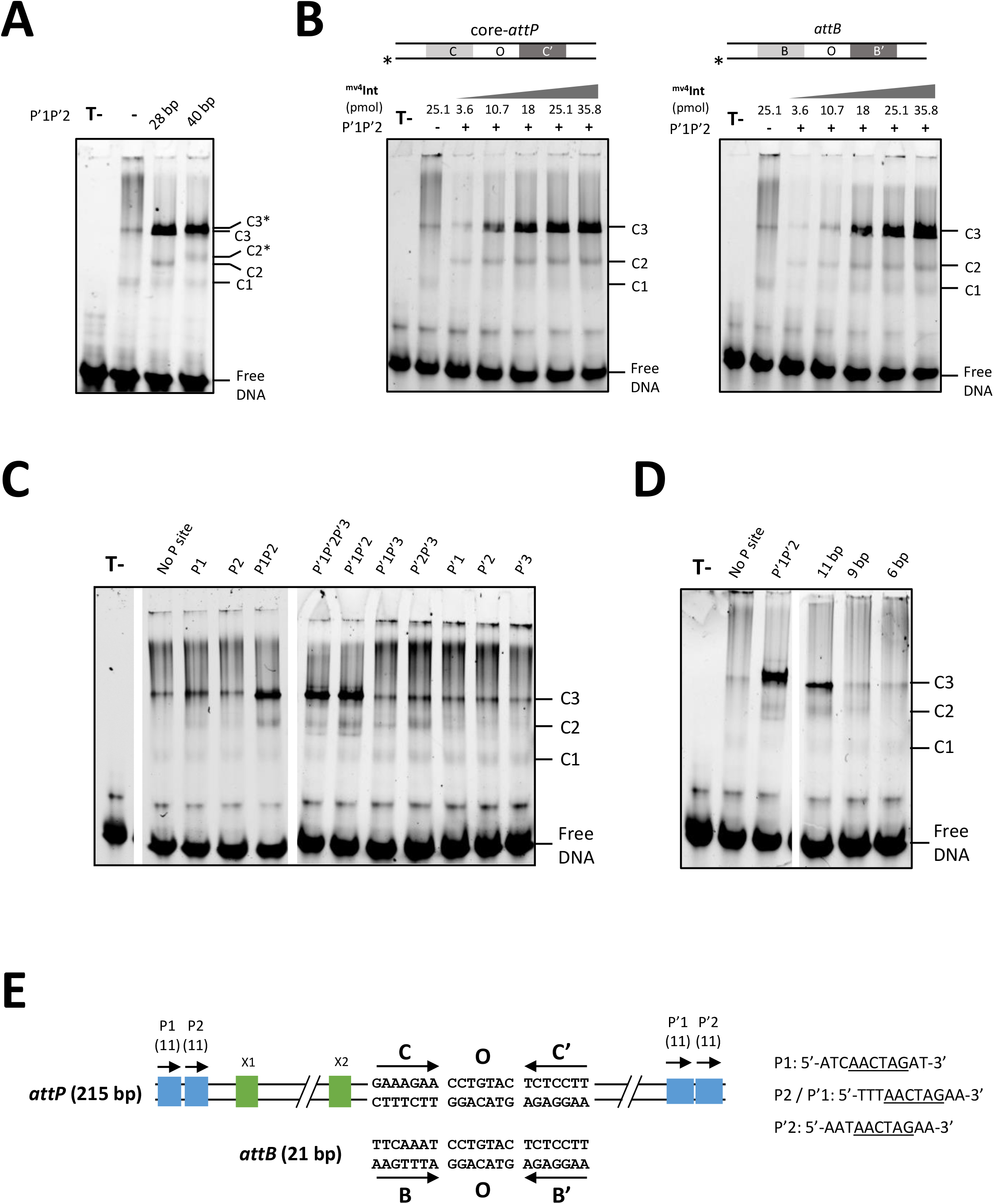
Characterisation of ^mv4^Int binding sites. (**A**) ^mv4^Int binding stabilisation to a 35-bp DNA fragment containing the core-*attP* region by different P’1P’2-containing unlabelled dsDNA fragments. EMSA reactions were performed as described in ‘Material and Methods’ with 25 pmol of ^mv4^Int. Presence/absence and size of arm sites are indicated above the gel. T-, control reactions (no ^mv4^Int, no arm site). Positions of the free DNA probes and the C1, C2/C2*, and C3/C3* ^mv4^Int/DNA complexes are indicated. (**B**) Titration of a 35-bp DNA fragment containing either the core-*attP* (COC’) or *attB* (BOB’) site by increasing concentrations of ^mv4^Int. The DNA substrate, ^mv4^Int concentrations in picomole, and presence/absence of P’1P’2 unlabelled DNA fragment are shown at the top. Free DNA as well as ^mv4^Int/DNA complexes are indicated. The asterisk locates the 3’-labelled strand. (**C**) Effect of 40-bp unlabelled dsDNA fragments containing different combination of arm sites on the ^mv4^Int binding stabilisation to the COC’ 35-bp core-*attP* fragment. Reactions and legends are identical to (**A**). (**D**) Size-dependence of the P’1P’2 site on the ^mv4^Int binding stabilisation to the 35-bp core-*attP* dsDNA. Reactions and legends are identical to (**A**). (**E**) New proposal for the genetic organisation of mv4 *attP* and *attB* recombination sites.

The arm-binding P sites were determined by assessing ^mv4^Int binding stabilisation to the WT core-*attP* (COC’) with several combinations of arm sites (Supplementary Table S3C). Two previous studies presented conflicting views on the size, location and orientation of the P sites (Supplementary Figure S6C). The first study proposed a 9-bp consensus site containing the conserved 6-bp 5’-AACTAG-3’ sequence, with P1, P2, P’1 and P’2 in direct repeats and P’3 in reverse orientation (39), whereas the second study suggested five11-bp sites all in direct repetition, with P2 adjacent to the core region (42). Among the eleven arm site combinations tested, only those containing either the adjacent P1P2 or P’1P’2 pairs effectively stabilized ^mv4^Int binding to COC’ (Figure 5C). This strongly indicated that the P’3 sequence is unlikely to serve as arm-binding site, confirming its unclear role in integrative and excisive recombination (39, 42). The size of the arm-binding sites was also evaluated by conducting binding experiments with 11-, 9-, or 6-bp P-site pairs (Supplementary Table S3C), revealing a stabilized complex C3 only with the two 11-bp sites in direct repeats (Figure 5D). Consequently, the *attP*_mv4_ site appears to be of 215 bp in length with two pairs of adjacent arm-binding sites in direct repeats, one pair (P1-P2) located at the end of the left arm, and the other (P’1-P’2) situated at the end of the right arm (Figure 5E).

### Highlighting compositional biases at the core-binding regions

The observed degeneracy within the mv4 *attB* and core-*attP* sites prompted us to hypothesize that ^mv4^Int could potentially catalyse site-specific recombination at alternative target sites, provided these targets follow the established consensus sequence. Recombination experiments using the *in vitro* fluorescent assay were conducted on several *attB* sites, modified to varying extents in their core-binding regions (Figure 6). These modified *attB* were tested against the WT *attP* site, or when *attB* overlap region was modified, against *attP* with overlap region identical to the modified *attB*. As expected, altering the B or B’ hemisite by substituting a nucleotide from the consensus sequence had no discernible effect on recombination (*attP*_4_/*B*_4_ and *attP*_5_/*B*_5_, Figure 6). Surprisingly, aside from the *attP*_1_/*B*_13_ pair, recombination remained effective even when one of the modified nucleotides was excluded or underrepresented in the *Sequence Logo* (*attB*_6_ to *attB*_10_, Figure 6). Impaired recombination only occurred when at least two “unfavourable” nucleotides were introduced, although it cannot be ruled out that recombination might have occurred with an efficiency below the detection limit of the method. These results provide an explanation for the erroneous determination of the minimal *attB* site previously reported (40). This *attB* site contained four nucleotides conforming to the B consensus sequence and only one “forbidden” nucleotide at the third position, which likely did not hinder recombination (*attP*_WT_/*attB*_min(99)_, Figure 6). In contrast, the shorter *attB* sites fortuitously contained two adjacent “unfavourable” nucleotides (*attP*_WT_/*attB*_3(99)_ and *attP*_WT_/*attB*_4(99)_, Figure 6), likely inhibiting recombination and leading the authors to mistakenly identify the left limit of the *attB* site at the penultimate nucleotide upstream the cleavage site. Given the lack of recombination observed for the *attP*_1_/*B*_13_ pair, despite containing only one unfavourable nucleotide, and assuming that *Sequence Logos* alone may not provide sufficient information to identify DNA motifs permissive to recombination or to determine the presence of other intrinsic constraints, we explored whether the enrichment ratio, calculated from recombination experiments with the random DNA libraries, could shed light on *in vitro* fluorescent assays. Assuming that the frequency of DNA motifs among permissive sites may, to some extent, correlate with their recombination efficiency, each random heptamer of the B, B’, C, and C’ DNA libraries was ranked based on its enrichment ratio (Supplementary File 2). These ranking values tended to highlight the recombination potential of hemisites, as the permissive pairs, except a*ttP*_6_/*B*_6_, contained both B’ and C’ hemisites with ranks higher than the wild-type. Conversely, the three nonpermissive pairs (*attP*_11_/*attB*_11_, *attP*_12_/*attB*_12_ and the 16-bp *attB*) contained B’ and C’ sequences negatively enriched or absent from the recombined site population (Figure 6). Ranking values partially explained the recombination deficiency of the *attP*_1_/*B*_13_ pair, as its B region, although containing nucleotides present in the consensus sequence, had a poorly ranked enrichment ratio (Figure 6). These ranking values also provided insights into intrinsic constraints on recombination sites. Except for the C site, which exhibited a low rank (2291) and enrichment ratio (2.4), all native core-binding sites were included in the “reduced” dataset of heptamers used to construct the *Sequence Logos* (Figure 4C), even if they were not necessarily among the top-ranked motifs (Table 1). The low values of one hemisite for *attP* (C) and one for *attB* (B’) may not be surprising, as they could be associated with evolutionary selection of recombination systems with low activity to minimize detrimental effect on the host, as exemplified for Tn*5* or Sleeping Beauty transposons where binding sites were found to be suboptimal for recombination (77, 78). Nonetheless, it cannot be ruled out that C and B’ hemisites are well adapted for *in vivo* but not *in vitro* ^mv4^Int-mediated recombination. The top-ranked motif for B’ and C’ hemisites contained the four central nucleotides (5’-CTCC-3’) seen on the wild-type sites, whereas B and C motifs retained only two or three nucleotides (Table 1). This asymmetrical conservation may be related to the enhanced binding capacity of B’/C’ sites, as complexes C2 and C3 were exclusively observed on these hemisites (Supplementary Figure S6B).

**Figure 6.**
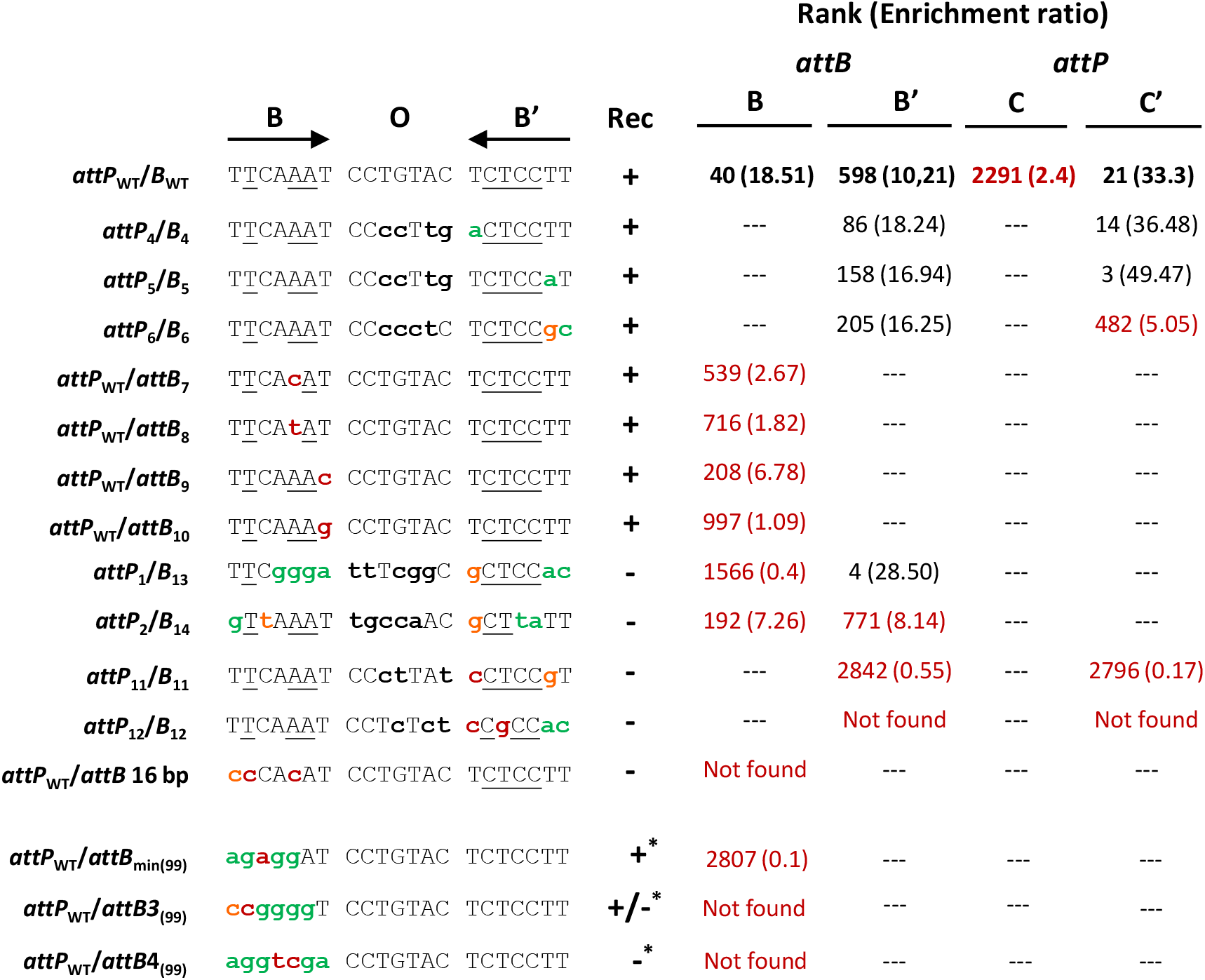
*In vitro* recombination outcomes of modified core-*attP* and *attB* sites. Nucleotides differing from the WT sites are represented by bold lowercase letters. Nucleotides present or absent from the *Sequence Logos* are indicated in green or red, respectively. Nucleotides underrepresented in the *Sequence Logos* are indicated in orange. +, recombination; +/-, recombination at low level; -, no recombination. Asterisk indicate *in vivo* recombination experiments described in (40). Rank and enrichment ratios are indicated for each hemisite. Values below the threshold used to determine “reduced” datasets are indicated in red. Not found: the DNA motif is absent from the recombined product population.

**Table 1.** Ranking values and enrichment ratios of remarkable motifs.

| Hemisite |  | DNA sequence | Rank (ratio) |
| --- | --- | --- | --- |
| B<br>(117 motifs) | Best | **a**<u>T</u>**gg**<u>AA</u>**a** | 1 (58.93) |
|  | Consensus | **g**<u>T</u>**gg**<u>AA</u>**a** | 2 (57.04) |
|  | WT | <u>TT</u><u>CAA</u><u>AT</u> | 40 (18.51) |
| B'<br>(613 motifs) | Best | **a**<u>CTCC</u>**c**T | 1 (46.53) |
|  | Consensus | **at**<u>T</u>**t**<u>C</u>TT | 136 (17.32) |
|  | WT | <u>TCTC</u>CTT | 598 (10.21) |
| C<br>(1025 motifs) | Best | **actga**<u>AA</u> | 1 (16.5) |
|  | Consensus | G**tgaa**AA | 181 (9.65) |
|  | WT | GAAAG<u>AA</u> | 2291 (2.44) |
| C'<br>(231 motifs) | Best | **a**<u>CTCC</u>**ac** | 1 (111.92) |
|  | Consensus | **at**<u>T</u>**t**<u>C</u>TT | 12 (35.38) |
|  | WT | <u>TCTC</u>CTT | 21 (33.31) |

Nucleotides differing from the wild-type sequences are in bold lowercase

Finally, several results have prompted us to consider the possibility that other compositional constraints, such as nucleotide interdependencies, may influence the structure of recombination sites. First, the recombination outcomes of several *attP*/*B* pairs (*attP*_WT_/*B*_7_ to *attP*_WT_/*B*_10_, and *attP*_2_/*B*_14_, Figure 6) were not fully explained by their ranking values. Second, the DNA motifs corresponding to the most frequently occurring nucleotides did not have the highest rank (Table 1). To explore this hypothesis statistically, we calculated Cramer’s V (53) on the “reduced datasets” of each core-binding site. This parameter quantifies the strength of association between two variables, and in the case of DNA motifs, it provides valuable insights into how a nucleotide can influence the nucleotide composition in its vicinity. Typically, Cramer’s V values above 0.2 indicate significant association. Varying degrees of nucleotide interdependencies were observed for many positions within the core-binding sites, with moderate to high interdependencies in seven or two pairs (B and B’, respectively) for the *attB* site, and two to three pairs (C and C’, respectively) for the core-*attP* site (Table 2). This seems to indicate a higher compositional bias on the bacterial genome site compared to the phage site. The most significant interdependencies were generally found within the central triplet, where moderate and strong associations were calculated for pairs 3-4, 4-5, and 3-5. An alternative approach to visualize nucleotide interdependency involved generating *Sequences Logos* for each hemisite while fixing one nucleotide at each position and assessing its impact on the distribution at other positions within the motif (Supplementary Figure S7). When this analysis was conducted on the WT hemisites, it confirmed the presence of interdependencies within the central 3-4-5 triplet for B and C hemisites, but not for B’ and C’ hemisites. Furthermore, it unveiled additional associations that had not detected using Cramer’s V (Supplementary Figure S7A). For instance, when the first position of the B hemisite was fixed to its WT nucleotide (T), it favoured the presence of an A at the sixth position. Similarly, fixing the last position to T increased the likelihood of observing two Gs at the third and fourth positions. Interestingly, when this analysis was conducted on the best heptamers (Table 1), the *Sequences Logos* revealed either the absence of interdependencies (B’ hemisite) or different interdependencies compared to those identified for the WT sites (Supplementary Figure S7B). In sum, these various analytical approaches collectively underscore the presence of multiple levels of constraint operating on the core-binding sites. This complexity suggests that establishing clear-cut criteria for predicting sequences efficient in ^mv4^Int-mediated recombination may be a challenging task.

**Table 2.** Nucleotide interdependencies on core-binding sites.

| Position 1 | Position 2 | Cramer's V before<br>recombination<br>(*attP* or *attB*) | Cramer's V after<br>recombination<br>(*attL* or *attR*) | Degree of freedom<br>(DoF) |
| --- | --- | --- | --- | --- |
| **hemisite B** |  |  |  |  |
| **3** | **4** | 0.020584 | **0.454310** | 2 |
| **3** | **5** | 0.008406 | **0.584629** | 1 |
| **4** | **5** | 0.020854 | **0.718980** | 1 |
| 4 | 6 | 0.007458 | 0.264725 | 2 |
| 5 | 6 | 0.025849 | *0.263922* | 1 |
| 5 | 7 | 0.011693 | *0.208535* | 1 |
| 6 | 7 | 0.030151 | 0.246502 | 2 |
| **hemisite B'** |  |  |  |  |
| 3 | 4 | 0.037191 | 0.284422 | 2 |
| 4 | 5 | 0.041570 | 0.249123 | 2 |
| **hemisite C** |  |  |  |  |
| 4 | 5 | 0.031626 | 0.204571 | 3 |
| 4 | 6 | 0.012116 | *0.165684* | 3 |
| **hemisite C'** |  |  |  |  |
| 1 | 5 | 0.008337 | *0.203884* | 2 |
| 2 | 3 | 0.020759 | *0.267007* | 1 |
| 3 | 4 | 0.018056 | 0.400487 | 1 |
| 3 | 5 | 0.013714 | 0.358316 | 1 |
| 4 | 5 | 0.021232 | 0.332503 | 2 |

Cramer's V values corresponding to high ( $V \geq 0.5$ for DoF 1, $\geq 0.35$ for DoF 2, and $\geq 0.29$ for DoF 3), moderate ( $V \geq 0.3$ for DoF 1, $\geq 0.21$ for DoF 2, and $\geq 0.17$ for DoF 3), and low ( $V \geq 0.1$ for DoF 1, $\geq 0.07$ for DoF 2, and $\geq 0.06$ for DoF 3) associations are indicated in **bold**, roman, and *grey italic*, respectively. Cramer's V background are calculated from the random libraries before recombination.

## CONCLUDING REMARKS

Previous investigations on the ^mv4^Int/*attP*/*attB* system have highlighted its uniqueness among heterobivalent integrases, notably i) the independence of species-specific host factor for recombination (39, 41), ii) the shortest *attB* site described in the literature, featuring an asymmetric organisation and the absence of core-binding sites (40), and iii) the peculiar architecture of ^mv4^Int arm-binding sites and the unusual size of its overlap region (42). Our study has provided a more comprehensive characterisation of this system, correcting and clarifying some of these properties, making the ^mv4^Int/*attP*/*attB* system the first fully characterized YR system of a temperate phage infecting a bacterial species within the *Bacillota* (ancient *Firmicutes*) phylum, and the second, after the mycophage L5 (34), outside the *Gammaproteobacteria*. ^mv4^Int has recently being phylogenetically classified as a member of the largest heterobivalent YRs family: the ^Tn916^Int subgroup, and more particularly within the streptococcal phage T12 cluster (25). The predicted structure of the corrected protein sequence confirms an organisation typical of heterobivalent YRs, and reveals the conserved β-hairpin insertion characteristic of the ^Tn*916*^Int subgroup. Structure-based alignment of the β-hairpin region from several YRs (Supplementary Figure S8A) revealed subtle variations in primary sequence as well as in length of the β-strands (4 to 6 aminoacids) and the loop (1 to 5 aminoacids). These variations resulted in two type of β-hairpin, “sharp” and “smooth” (Supplementary Figure S8B), which could differ in their interaction with the recombination sites. ^mv4^Int and its relatives seem to contain longer β-strands (6-aa) separated by only one aminoacid, producing a sharper structure compared to the representative ^T12^Int or other integrases from the ^Tn916^Int subgroup such as ^Tn*1549*^Int or ^CTn6^Int. Whether this structural difference has any functional consequence regarding the role of the β-hairpin requires further experimental investigation. We confirmed that ^mv4^Int does not require species-specific host factor to recombine WT *attP* and *attB* sites. However, we demonstrated that the ubiquitous NAP HU, while not mandatory, can significantly enhance the recombination between *attP*/*B* sites, particularly the modified ones. A similar stimulation has been observed for Tn*916* excision in *E. coli* (79). HU is frequently found as cofactor for serine recombinases, such as Tn*3*/γδ or Hin, (80–83), but is relatively rare in model heterobivalent YRs, which typically use IHF (27, 29, 84, 85) or analogs like mIHF for ^L5^Int (86) or BHFa for ^CTnDOT^Int (87). HU and IHF belong to the same DNA bending family, with IHF being exclusively found in *Proteobacteria* (88, 89). They differ in their substrate preference and bending ability (90). It’s worth noting that, except for ^L5^Int, all members of the ^Tn916^Int subgroup studied to date (91–93) are functional on different bacterial species, confirming their independence from host factors, similar to ^mv4^Int. We hypothesise that the β-hairpin found in ^Tn916^Int subgroup, through its bending activity (57), shapes the *attP* site to facilitate intasome formation with the assistance of HU, in contrast to YRs lacking the β-hairpin which may require more specific host factor with strong bending activity, such as IHF or BHFa, to form the intasome.

The use of *in vitro* recombination involving randomized DNA libraries, PCR amplification of recombination products, and subsequent Sanger sequencing or NGS analysis, has proven to be an effective approach in characterising the core regions *attB* and *attP* sites. This methodology appears well-suited for the investigation of recombination sites and has been successfully employed in various recombination systems (59, 71, 94, 95). However, it is important to note that in the context of site-specific recombination systems, obtaining PCR products from recombined junctions (*attL* or *attR*) does not necessarily guarantee a complete reaction, i.e. the two successive strand-exchange, as it is possible to obtain PCR amplicons as soon as the first strand-exchange occurred and generated the Holliday junction (HJ). Nevertheless, we considered the presence of *attL* or *attR* PCR fragments as representative of site-specific recombination, because several YRs systems, such as CTXϕ (96), VGJϕ (97) or integrons (98, 99), are defined as site-specific recombination systems while only the first strand-exchange is catalysed by the recombinase, with the HJ being subsequently resolved by the host replication machinery. This experimental strategy led us to demonstrate that mv4 *attB* site is longer than previously described, with a size of 21 bp more consistent with other heterobivalent YRs systems ranging from 18 bp for HP1 phage (63) to 29 bp for the mycophage L5 (64). This approach unequivocally characterised the size of the overlap region as a 7-bp sequence, the canonical size for heterobivalent YRs systems found in bacteriophages, and revealed no constraint on nucleotide composition as long as strict identity between *attB* and *attP* regions is maintained. However, when several overlap regions were tested by classical *in vitro* fluorescent assays, a significant effect of the sequence on recombination efficiency was observed, suggesting compositional biases to some extent. This particular aspect deserves further investigation through NGS analysis of the recombined products from the two *attB* × *attP* libraries containing randomized overlap regions. Furthermore, the randomized DNA libraries revealed a high level of nucleotide degeneracy on both 7-bp regions surrounding the overlap region of *attB* and core-*attP* sites. This nucleotide degeneracy led to the determination of consensus sequences that were nearly identical on both *attB* and core-*attP* site. As these left- and right sequences were complementary to each other, defining two inverted-repeated regions, this allowed us to propose an organisation of the mv4 recombination sites typical of the YRs systems, BOB’ on *attB* and COC’ on core-*attP*, with B, C, and B’/C’ hemisites corresponding to the genuine core-binding sites for ^mv4^Int. The level of nucleotide degeneracy observed on these core regions underscores a tolerance of ^mv4^Int to nucleotide variation that is generally unexpected for YRs, although very few studies focused on modifications of the core-binding (59, 68, 94, 100–102). Analysis of secondary integration sites of phage λ showed that despite nucleotide changes at B and B’ sites, a majority of the secondary sites contained the inverted-repeated sequence, indicating that ^λ^Int is capable of binding sequences other than the native B and B’ sequences, albeit with limited flexibility since both sequences must possess the 5’-CTT-3’ triplet (100). A similar requirement was observed for ^HK022^Int (68, 102). As our NGS data revealed no invariant nucleotide at any position of B and B’ sequences, this strengthens our hypothesis of a greater flexibility of ^mv4^Int towards its recombination sites. Therefore, it would be interesting to use the randomized DNA libraries methodology to other model YR systems to investigate whether the flexibility observed in ^mv4^Int is an intrinsic characteristic of this protein or if it is a more common property among heterobivalent integrases. Additionally, since phage mv4 belongs to class III of temperate phages, which integrate their genome at the 3’ end of tRNA gene (103), utilising asymmetric *attB* site lacking IR sequences (104), it would be valuable to explore if flexibility is more closely related to this particular group of integrases.

Finally, our exploration of the consensus sequences derived from the random DNA libraries, coupled with classical gel-based recombination assays, has shed light on the intricate interplay of various compositional constraints governing the adaptation of recombination sites to their cognate integrase. These constraints, which exert cumulative effects, challenge the predictability of recombination efficiency when modifying such sites. In addition, it is important to note that our recombination experiments involved only one randomized core-binding site at a time, either left or right, thereby potentially overlooking additive effects that may occur between the two sites. Nonetheless, we envision harnessing the ^mv4^Int flexibility towards its recombination sites to develop a programmable genetic tool by adapting the *attP* site to novel DNA targets. This will require further developments, such as enhancing recombination efficiency by testing *attP* sites with highly-ranked core-binding motifs. Additionally, conducting *in vivo* recombination experiments on bacterial species of biotechnological interest, using sites with randomized core-*attP* regions, will aid in identifying integration sites that could serve as future “landing pads”, as recombination reactions appear to be more permissive *in vivo* than *in vitro* in some recombination systems (59, 95). In summary, this study underscores the diversity and complexity of recombination systems employed by temperate bacteriophages. It offers valuable insights into the structure and functioning of the ^mv4^Int/*attP*/*attB* recombination module, thus expanding our comprehension of the mechanisms underpinning site-specific recombination and the determinants contributing to the biological diversity of these systems.

## SUPPLEMENTARY DATA

1 pdf supplementary data + 1 xlsx heptamer ranking

Supplementary Data are available at NAR online.

## Supporting information

Supplementary file 2_heptamers enrichment ranking

## ACKNOWLEDGEMENT

We thank Julie De Stefano, Clementine Duarte, and Romain Bernardo for technical assistance. We thank Céline Loot for helpful discussion during the writing of the manuscript.

## Author contributions

K.D., and P.L.B. designed the project and wrote the original draft of the manuscript; K.D., P.L. and M.P. performed the experiments; M. Campos performed the bioinformatic analysis; K.D., M. Coddeville, M. Campos, P.R., M.C.B. and P.L.B. analysed the data; All authors contributed to reviewing and editing.

---

**Supplementary Figure S1.**
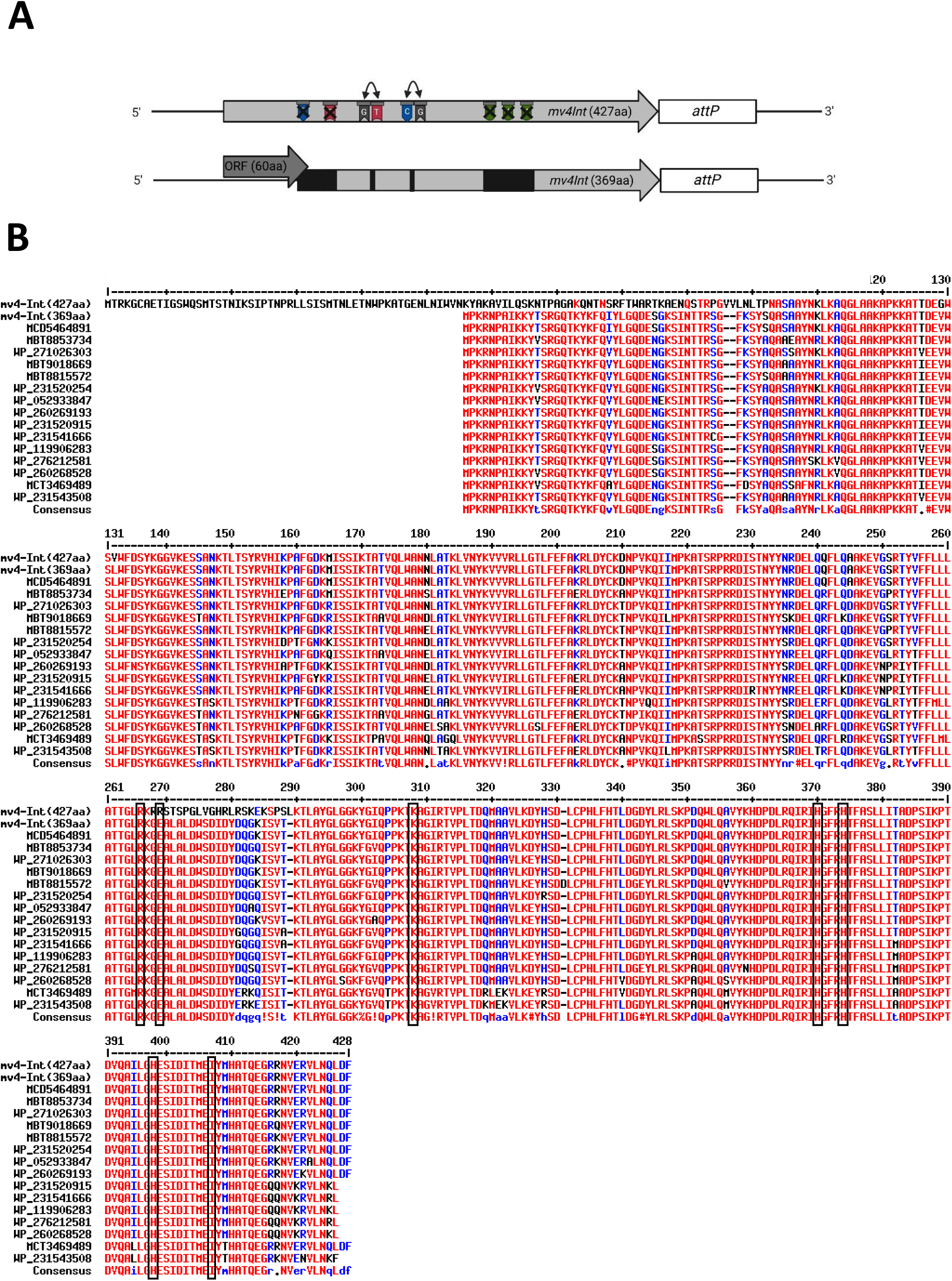
(**A**) **Schematic representation of ^mv4^Int(427aa) encoding gene and consequences of sequencing errors.** Three deletions (one C, one T and three A; black crosses) and two inversions (GT/TG and CG/GC; two-way arrows) were found. Black boxes indicate protein regions differing from the published sequence (accession number U15564). (**B**) **Multiple sequence alignment of *L. delbrueckii* subsp. *bulgaricus* prophages integrase phylogenetically close to ^mv4^Int.** A BLAST search was performed against the ^mv4^Int369 protein sequence. Sequences with identity higher than 80% were collected and subjected to alignment using Multalign algorithm (Corpet, 1988). The two first sequences correspond to the published ^mv4^Int (Dupont *et al.*, 1995) and the sequence obtained from this study, respectively. MCD5464891 *L. delbrueckii* subsp. *bulgaricus* strain CIRM BIA 773 (identical to MCD5482372, *L. delbrueckii* subsp. *bulgaricus* strain CIRM BIA 2159); MBT8853734, hypothetical protein BTH55_02920 *L. delbrueckii* subsp. *bulgaricus* strain XJ77-7-2; WP_271026303, site-specific integrase *L. delbrueckii*; MBT9018669, hypothetical protein BTI91_04510 *L. delbrueckii* subsp. *bulgaricus* strain MGC24-1; MBT8815572, hypothetical protein BTH97_01860 *L. delbrueckii* subsp. *bulgaricus* strain XJ13-13; WP_231520254, site-specific integrase *L. delbrueckii*; WP_052933847, site-specific integrase *L. delbrueckii*; WP_260269193, site-specific integrase *L. delbrueckii*; WP_231520915, site-specific integrase *L. delbrueckii*; WP_231541666, specific integrase *L. delbrueckii*; WP_119906283.1, site-specific integrase *L. delbrueckii*; WP_276212581, specific integrase *L. delbrueckii*; WP_260268528 specific integrase *L. delbrueckii*; MCT3469489, site-specific integrase *L. delbrueckii* subsp. *bulgaricus* CIRM-BIA 1579; WP_231543508, specific integrase *L. delbrueckii.* High (>90%) and low (>50%) consensus values are indicated in red and blue, respectively. Variable residues are indicates in black. The seven conserved residues of the catalytic pocket (Gibb *et al.*, 2010) are boxed.

**Supplementary Figure S2.**
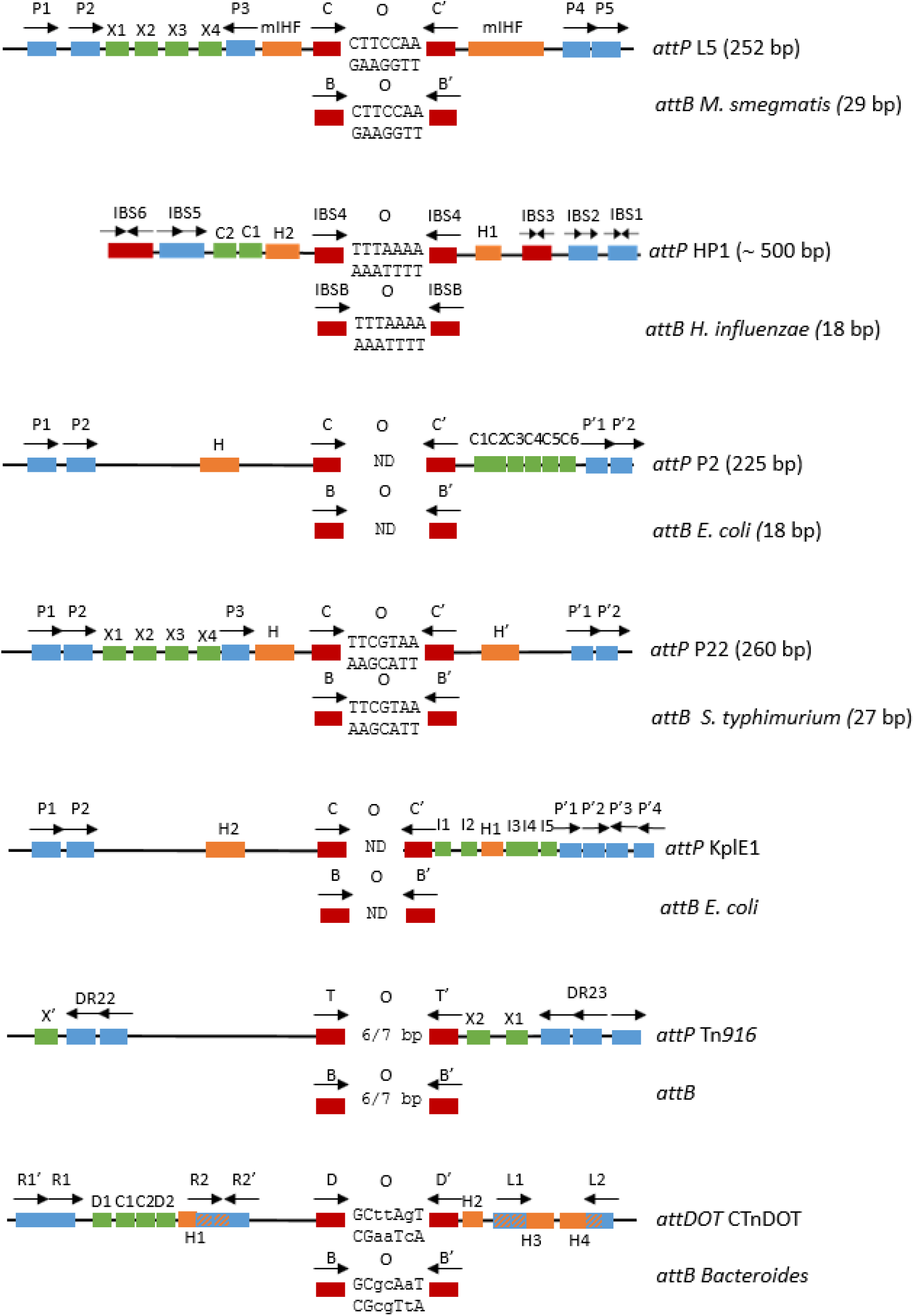
Genetic organisation of L5, HP1, P2, P22, KlpE1, Tn916, and CTnDOT recombination sites. Arm-binding sites are represented by blue rectangles. Core-binding sites are represented by red rectangles. Orange rectangles indicate the binding sites of the host-factors. The black arrows indicate the orientation of the binding sites. In the case of CTnDOT, the hatching indicates the overlap of an IHF-binding site with an arm-binding site. Green rectangles indicate the excisionase binding sites (Xis for λ, L5, P22, Tn916, mv4, Xis2c and Xis2d for CTnDOT, Cox for HP1, P2 and TorI for KplE1). The sequences of the central regions are represented, and lowercase letters indicate the nucleotides of the overlap region that are different between the *att*DOT and *attB*.

**Supplementary Figure S3.**
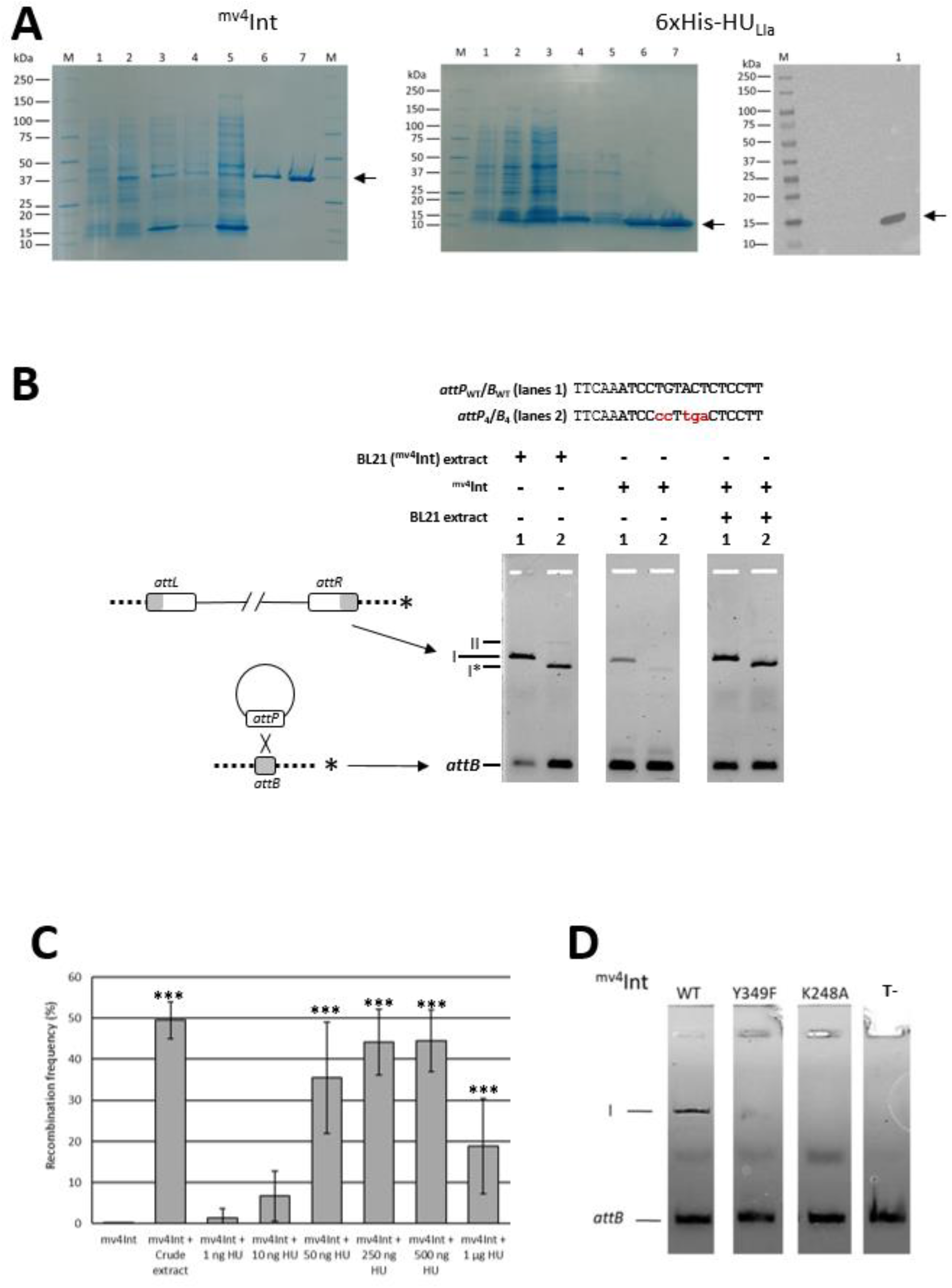
(**A**) **Protein purification.** Left, ^mv4^Int purification. SDS-PAGE: Lanes: 1, uninduced *E. coli* BL21(DE3) crude extract; 2, induced crude-extract; 3, soluble fraction; 4, insoluble fraction; 5, Ni-NTA column flow-through; 6, gel filtration eluted fractions; 7, Heparin column eluted fractions; M, protein ladder. Right, 6xHis-HU_Lla_ purification. SDS-PAGE: Lanes: 1, uninduced *E. coli* BL21(DE3) crude extract; 2, induced crude-extract; 3, supernatant before heat treatment (70°C, 30 min); 4, supernatant after heat treatment; 5, Ni-NTA column flow-through; 6, Ni-NTA column eluted fractions; 7, Heparin column eluted fractions; M, protein ladder. Western-blot: 200 ng of 6xHis-HU_Lla_ were separated by SDS-PAGE, transferred to a nitrocellulose membrane and detected by chemiluminescence after incubation with Anti-polyHistidine−Peroxidase (1: 2000). Lanes: 1, 200 ng 6xHis-HU_Lla_; M, protein ladder. (**B**) ***In vitro* recombination assays** with *E. coli* BL21 overexpressing ^mv4^Int crude extract or purified ^mv4^Int on WT (lanes 1) or modified sites (lanes 2). The 16-bp minimal *attB* site (Auvray *et al.*, 1999b) is indicated in bold. Nucleotides differing from the WT sites are indicated in red lowercase letters. Linear fluorescent *attB* fragment and recombination products between wild-type sites (I) or modified sites (I*) are indicated. The second recombination product (II) is most likely a secondary recombination product from an *attP* x *attL* recombination (Coddeville and Ritzenthaler, 2010). Size difference between recombined fragments I and I* is due to size difference of the plasmid carrying the *attP* sites, with the plasmid containing the *attP*_WT_ (pMET311) being 1.2-kb larger than the plasmid containing modified *attP* site (pMET306). (**C**) **Dose-dependence of purified 6xhis-HU_Lla_ protein** on recombination efficiency of the modified *attP*/*B* sites. Fluorescence intensity was quantified with Image Lab software (Biorad). The data presented are the mean values and standard deviations of four independent measurements. Significant differences from the ^mv4^Int condition are represented by the symbol “***” (Kruskal-Wallis p<0.001). **(D**) ***In vitro* recombination activity of ^mv4^Int variants**. Protein variant is indicated above each lane. T-, control reactions (no ^mv4^Int). The fluorescent 290-bp *attB* fragment and the linear recombination product (I) are indicated.

**Supplementary Figure S4.**
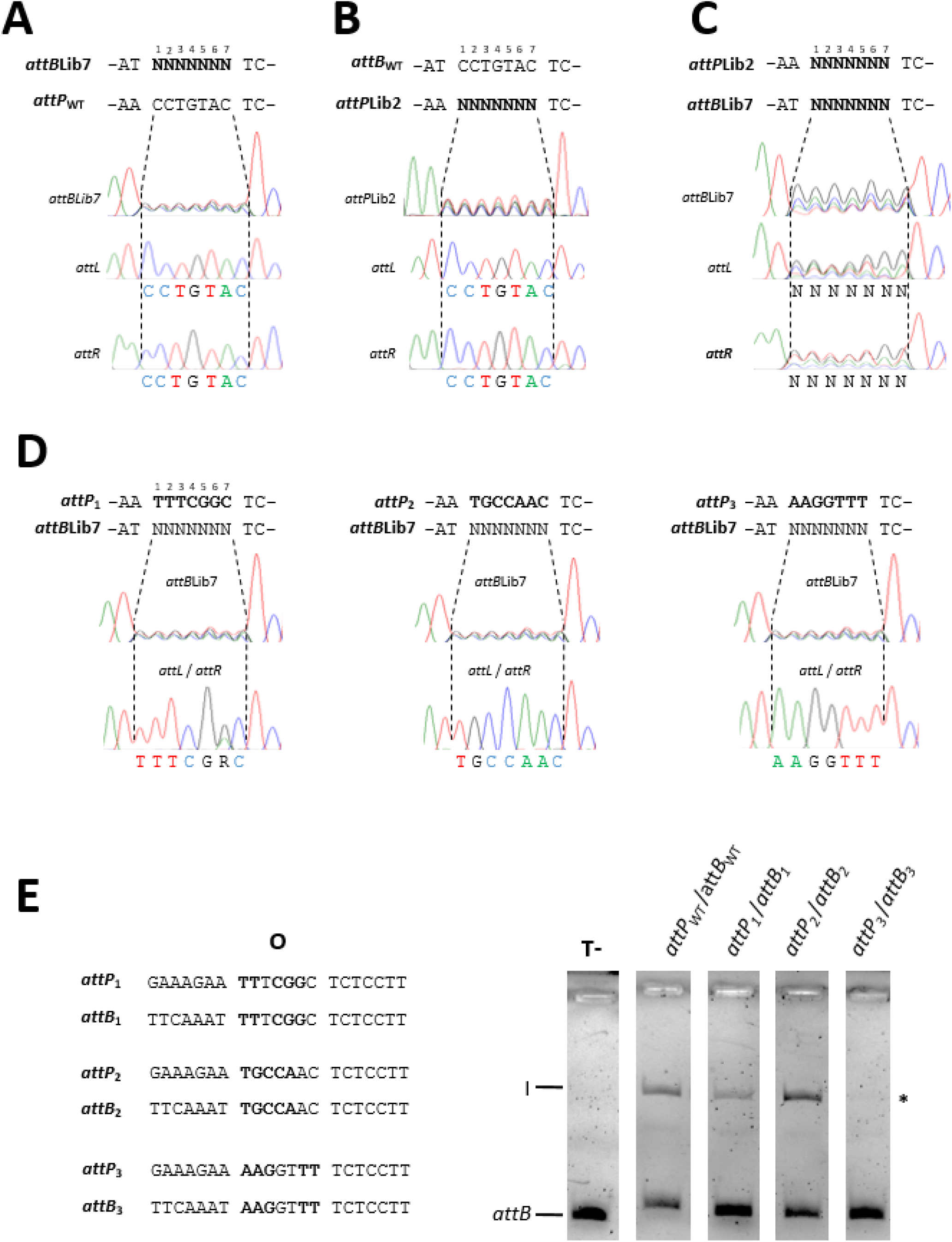
Constraints exerted at the strand-exchange region. (**A to C**) Chromatograms before (*attB* or *attP*) or after (*attL, attR*) recombination from *attB*Lib7 x *attP*_WT_ (n=3), *attB*_WT_ x *attP*Lib2 (n=2), and *attB*Lib7 x *attP*Lib2 *in vitro* reactions (n=2). **(D)** Chromatograms of *attB*, *attL, attR* regions of *attB*Lib7 recombination against three *attP* sites with different overlap regions. (**E**) *In vitro* recombination assay of the *attP*/*attB* pairs with the three overlap regions. Reactions were performed as described in ‘Material and Methods’. The fluorescent 290-bp *attB* fragment and the linear recombination product (I) are indicated. T-, control reactions (no ^mv4^Int). Asterisk indicates recombination product at the detection limit of the method.

**Supplementary Figure S5.**
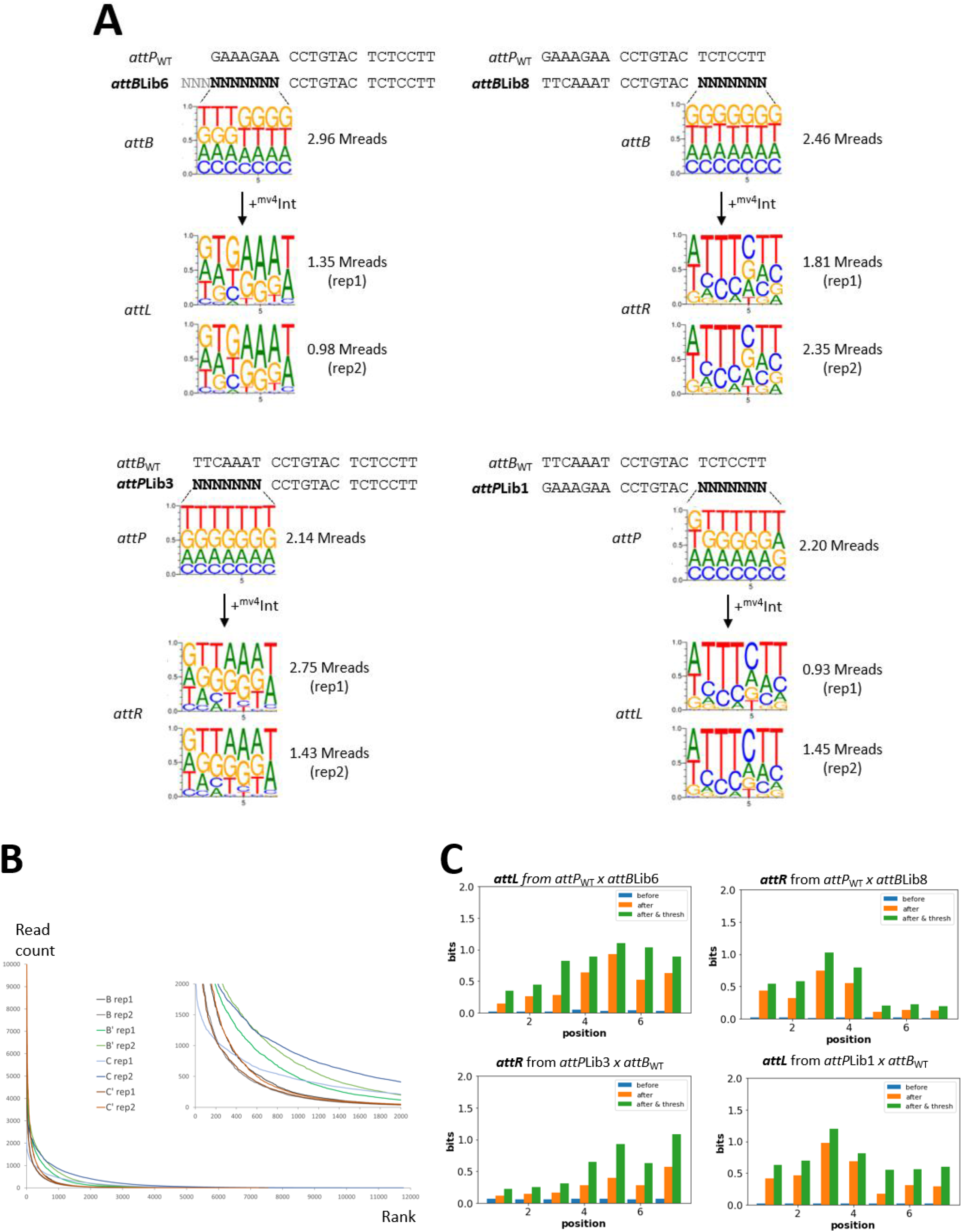
(**A**) *Sequence Logos* showing the nearly random nucleotide composition of *attP* and *attB* DNA libraries and the composition biases of *attL* or *attR* after recombination. Number of sequencing reads is indicated from each experiment. Rep1 and rep2 correspond to biological replicate. (**B**) Occurrence (read count) of each randomized heptamer relative to its ranking distribution on raw NGS data after recombination. B, *attL* region from *attP*_WT_ x *attB*Lib6 recombination; B’, *attR* region from *attP*_WT_ x *attB*Lib8 recombination; C, *attR* region from *attB*_WT_ x *attP*Lib3 recombination; C’, *attL* region from *attB*_WT_ x *attP*Lib1 recombination. (**C**) Factors affecting normalization of nucleotide prevalence at each position of the heptamers. Mutual information (MI) was evaluated using the following formula 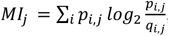, where *p_i,j_*(*q_i,j_*) is the probability of nucleotide *i* at position j in the enriched (background) motif. The distance value (bits) reflect the difference of nucleotide frequency between the ’observed’ and ’hypothetical’ random distribution. A value of 0 indicates no difference between observed and theoretical frequency for a random distribution. The maximal distance value is 2 (given by the formula *MI_j_max* = 4 * 0.25 * *log*_2_0.25, equal frequency of each of the 4 nucleotides). Before recombination, the initial set of sequences exhibits a near-random distribution, even after being weighted by the read counts (blue bars). After recombination, specific nucleotides are enriched at each of the seven positions by applying read count-based weighting (orange bars), and the same pattern is enhanced when applying a threshold based on the enrichment factor before and after recombination (green bars, below).

**Supplementary Figure S6.**
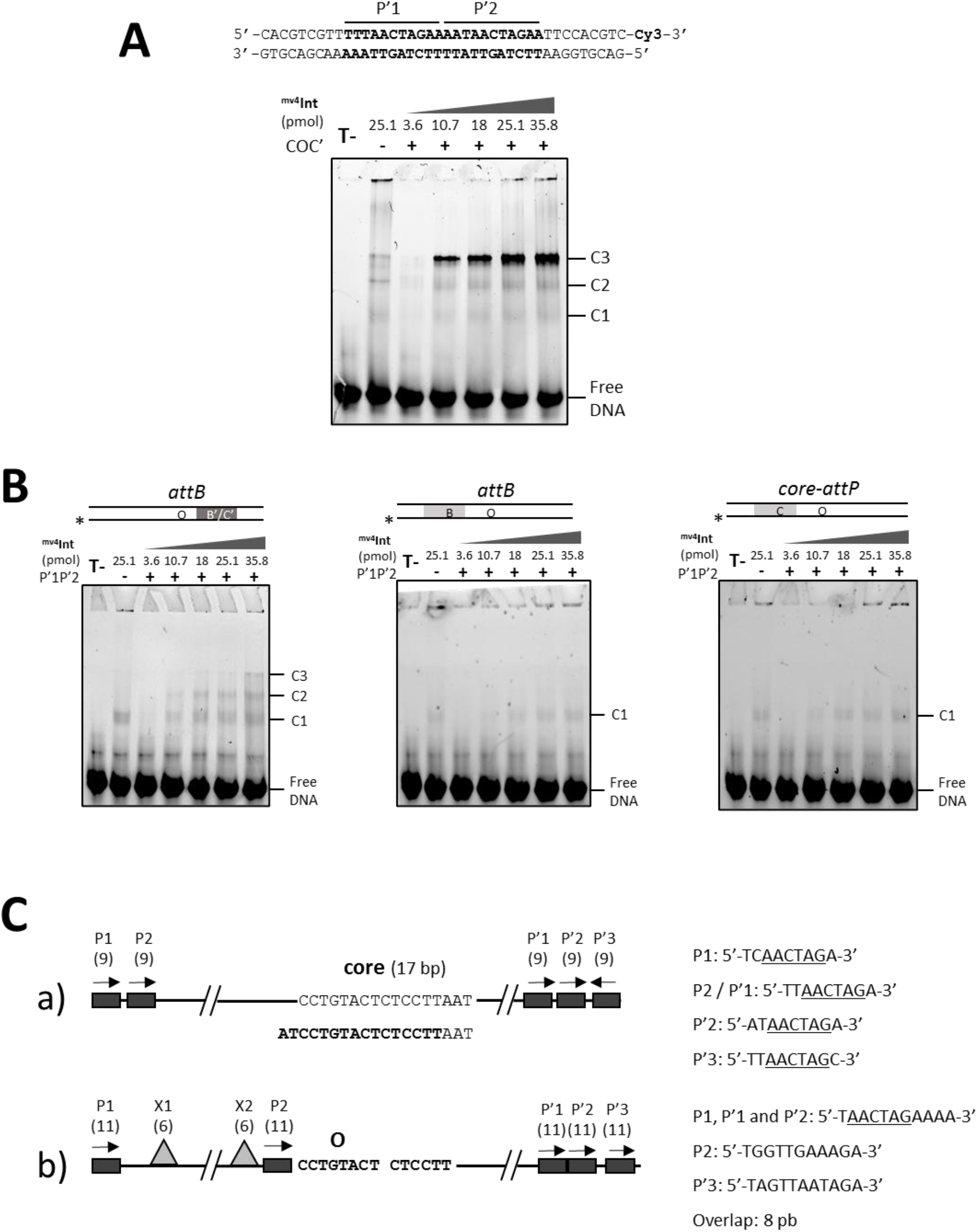
(**A**) Titration of a 40-bp DNA fragment containing the P’1P’2 sites by core-*attP* fragment with increasing concentrations of ^mv4^Int. The DNA substrate, ^mv4^Int concentrations, and presence/absence of unlabelled COC’ fragment are shown at the top. Free DNA and ^mv4^Int /DNA complexes are indicated. T-, control reactions (no ^mv4^Int, no core-*attP* site). (**B**). EMSA titration of a 35-bp *attB* or core-*attP* DNA fragment containing mutated left- or right hemisites. The DNA substrate is shown at the top, ^mv4^Int concentrations are given in picomole. Free DNA and ^mv4^Int/DNA complexes are indicated. The asterisk locates the 3’-labelled strand. (**C**) The different organisations of mv4 *attP*/*attB* recombination sites. Nucleotide sequence of P sites are indicated. The 16-bp minimal *attB* site is indicated in bold. a) initial description of the recombination sites (Dupont *et al.*, 1995; Auvray *et al.*, 1999a; Auvray *et al.*, 1999b); b) alternative P-arm sites organisation, characterization of Xis binding sites (grey triangle), and size of overlap region (Coddeville and Ritzenthaler, 2010; Coddeville *et al.*, 2014).

**Supplementary Figure S7.**
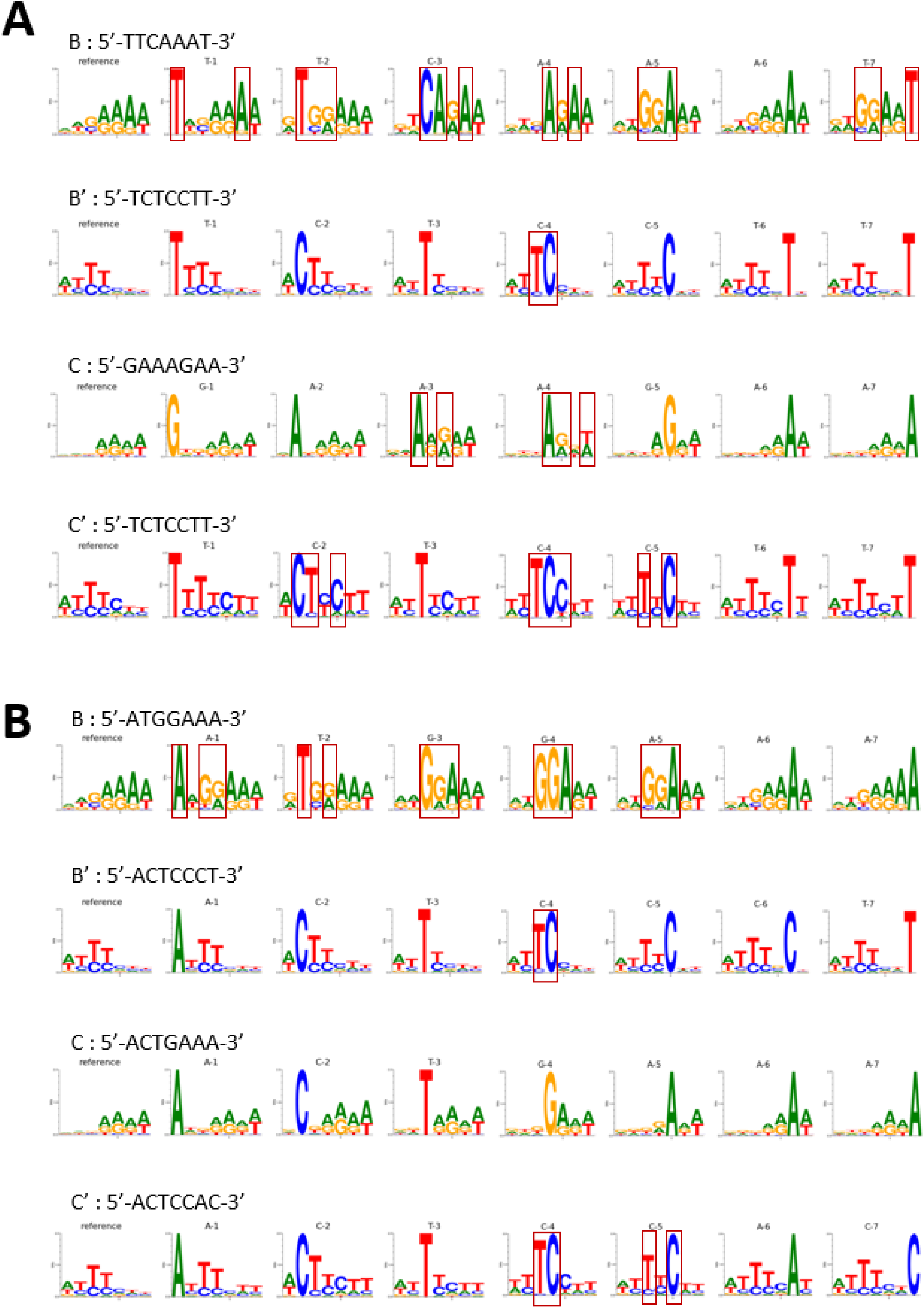
*Sequence Logos* of the *attP* and *attB* core-binding. (**A**) Fixing each nucleotide from the WT hemisites. (**B**) Fixing each nucleotide from the top-ranked hemisites.

**Supplementary Figure S8.**
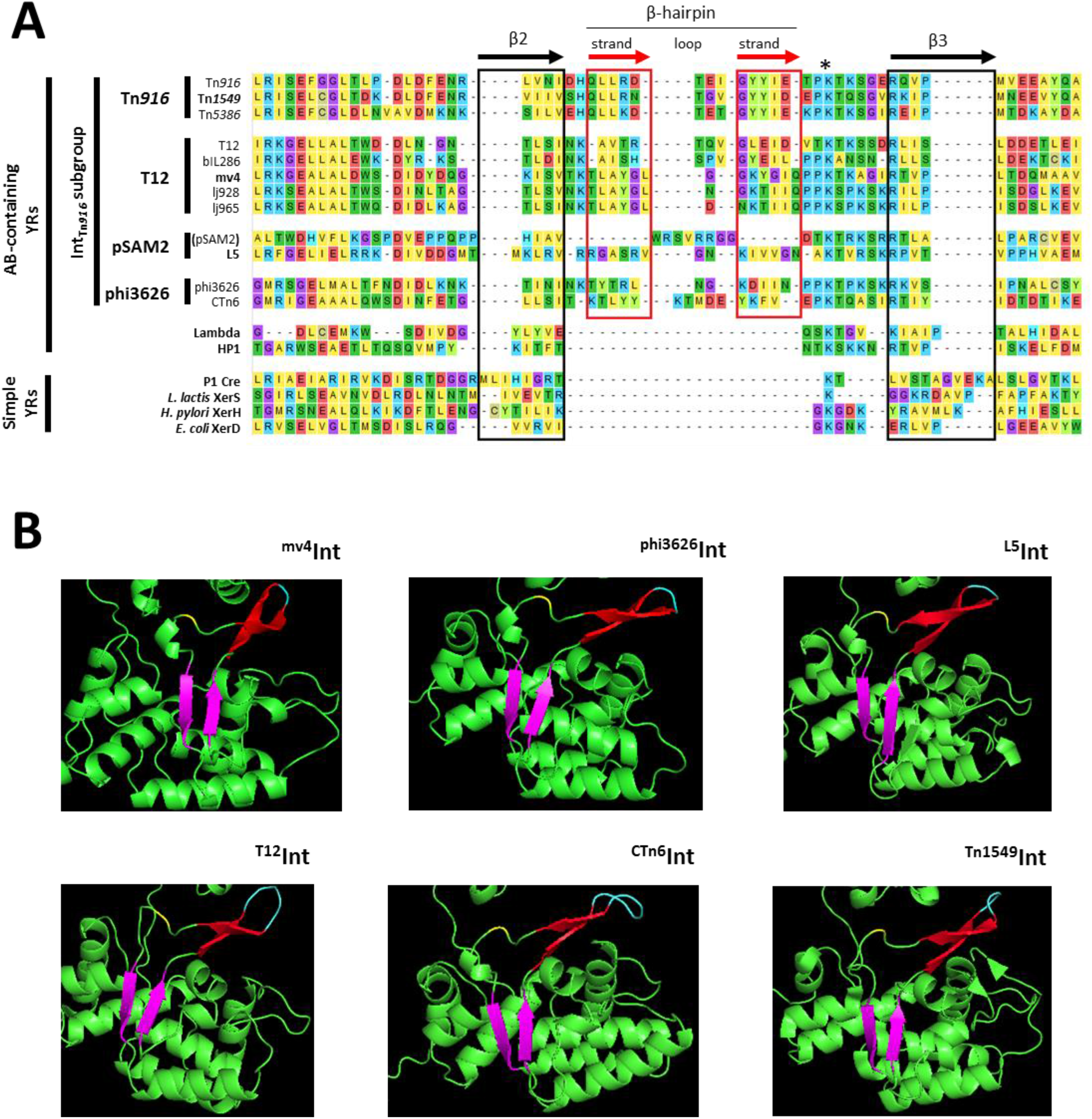
(**A**) Structure-based alignment of the β2-β3 region of Int_Tn916_ subgroup and other model Y-recombinases. Limits of β-strands were determined from X-ray data or Alphafold2 predictions. YR are classified according to Smyshlyaev *et al.*, 2021. The β-strands present in all YR are named according to Aihara *et al.*, 2003 and boxed in black. β-strands of the additional β-hairpin are boxed in red. The catalytic K residue (corresponding to K235 for ^λ^Int) is indicated by an asterisk. (pSAM2) : no β-hairpin could be observed for ^pSAM2^Int. (**B**) Details of the β2-β3 region revealing the two types of β-hairpins. Upper, “sharp” hairpins with 1 or 2 aminoacids on the loop; Lower, “smooth” hairpins with 3 to 5 aminoacids on the loop. β2-β3 strands are in purple. β-strands from the β-hairpin are in red, and loop in cyan. The catalytic K residue is in yellow.

**Table S1.** Bacterial strains.

| Name | Relevant properties | Reference |
| --- | --- | --- |
| *E. coli*<br>BL21(DE3) | B F<sup>-</sup> *ompT gal dcm lon hsdSB*(<sub>RB-MB-</sub>) $\lambda$(DE3 [*lacI lacUV5-T7p07 ind1 sam7 nin5*]) [*malB*<sup>+</sup>]<sub>K-12</sub>($\lambda^S$) | (Studier *et al.*, 1990) |
| *E. coli*<br>EPI300™ | F<sup>-</sup> $\lambda^-$ *recA1 endA1 mcrA* $\Delta$(*mrr-hsdRMS-mcrBC*) Φ80d[Δ*lacZ58*(M15) $\Delta$(*lac*)X74 *araD139* $\Delta$(*ara, leu*)7697 *galU galK rpsL* (Str<sup>R</sup>) *nupG trfA dhfr* | Lucigen |
| *E. coli*<br>MET961 | NEB 5-alpha *glgB::repA*(pWV01), KanR. Used for the propagation, cloning or modification steps of pMET340 and derivatives | Laboratory collection |

**Table S2.**
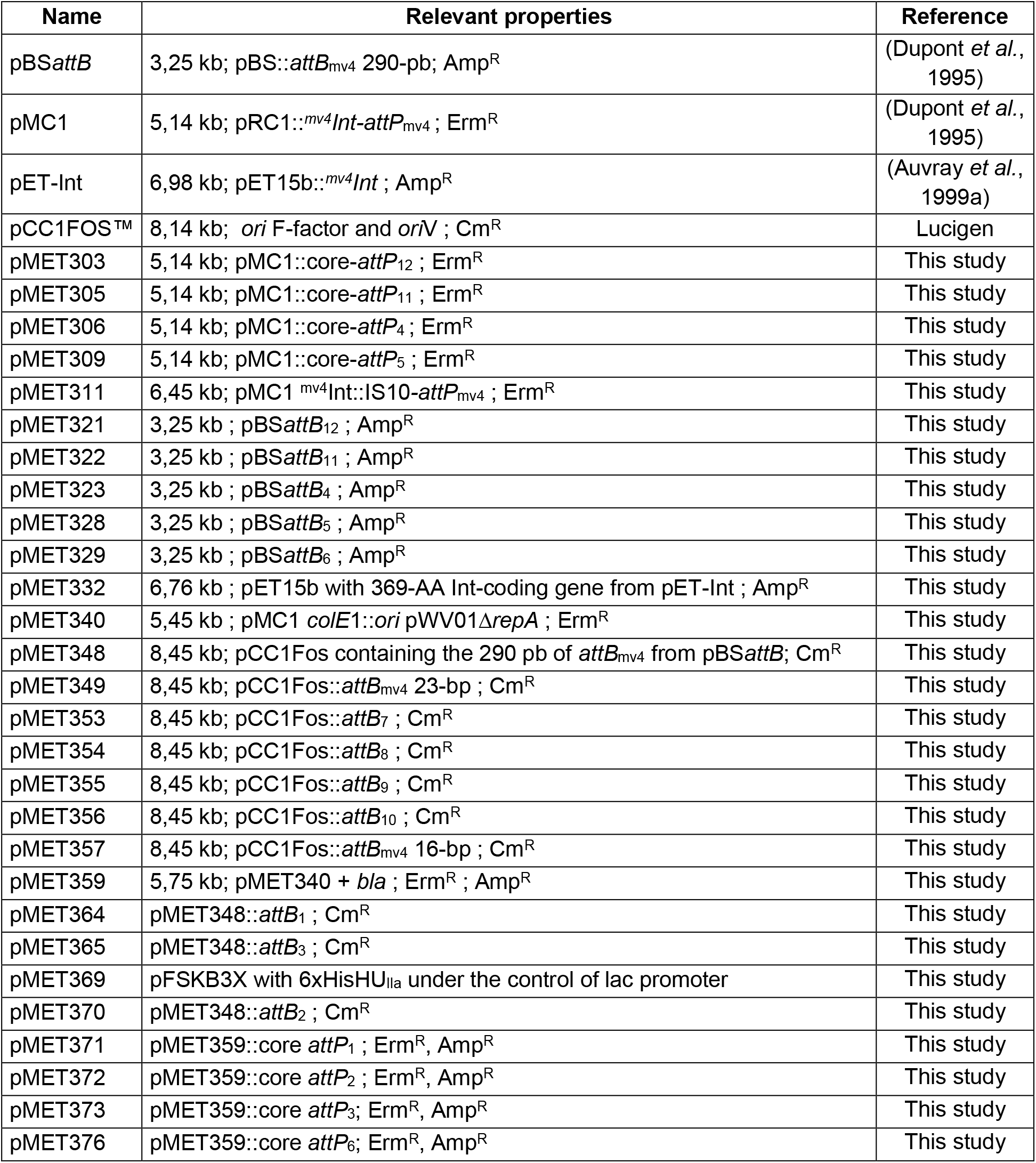
Plasmids.

| Name | Relevant properties | Reference |
| --- | --- | --- |
| pBSattB | 3,25 kb; pBS::*attB*<sub>mv4</sub> 290-pb; Amp<sup>R</sup> | (Dupont *et al.*, 1995) |
| pMC1 | 5,14 kb; pRC1::*mv4Int-attP*<sub>mv4</sub>; Erm<sup>R</sup> | (Dupont *et al.*, 1995) |
| pET-Int | 6,98 kb; pET15b::*mv4Int*; Amp<sup>R</sup> | (Auvray *et al.*, 1999a) |
| pCC1FOS™ | 8,14 kb; *ori* F-factor and *oriV*; Cm<sup>R</sup> | Lucigen |
| pMET303 | 5,14 kb; pMC1::*core-attP*<sub>12</sub>; Erm<sup>R</sup> | This study |
| pMET305 | 5,14 kb; pMC1::*core-attP*<sub>11</sub>; Erm<sup>R</sup> | This study |
| pMET306 | 5,14 kb; pMC1::*core-attP*<sub>4</sub>; Erm<sup>R</sup> | This study |
| pMET309 | 5,14 kb; pMC1::*core-attP*<sub>5</sub>; Erm<sup>R</sup> | This study |
| pMET311 | 6,45 kb; pMC1 <sup>mv4</sup>Int::IS10-*attP*<sub>mv4</sub>; Erm<sup>R</sup> | This study |
| pMET321 | 3,25 kb; pBSattB<sub>12</sub>; Amp<sup>R</sup> | This study |
| pMET322 | 3,25 kb; pBSattB<sub>11</sub>; Amp<sup>R</sup> | This study |
| pMET323 | 3,25 kb; pBSattB<sub>4</sub>; Amp<sup>R</sup> | This study |
| pMET328 | 3,25 kb; pBSattB<sub>5</sub>; Amp<sup>R</sup> | This study |
| pMET329 | 3,25 kb; pBSattB<sub>6</sub>; Amp<sup>R</sup> | This study |
| pMET332 | 6,76 kb; pET15b with 369-AA Int-coding gene from pET-Int; Amp<sup>R</sup> | This study |
| pMET340 | 5,45 kb; pMC1 *colE1::ori* pWV01Δ*repA*; Erm<sup>R</sup> | This study |
| pMET348 | 8,45 kb; pCC1Fos containing the 290 pb of *attB*<sub>mv4</sub> from pBSattB; Cm<sup>R</sup> | This study |
| pMET349 | 8,45 kb; pCC1Fos::*attB*<sub>mv4</sub> 23-bp; Cm<sup>R</sup> | This study |
| pMET353 | 8,45 kb; pCC1Fos::*attB*<sub>7</sub>; Cm<sup>R</sup> | This study |
| pMET354 | 8,45 kb; pCC1Fos::*attB*<sub>8</sub>; Cm<sup>R</sup> | This study |
| pMET355 | 8,45 kb; pCC1Fos::*attB*<sub>9</sub>; Cm<sup>R</sup> | This study |
| pMET356 | 8,45 kb; pCC1Fos::*attB*<sub>10</sub>; Cm<sup>R</sup> | This study |
| pMET357 | 8,45 kb; pCC1Fos::*attB*<sub>mv4</sub> 16-bp; Cm<sup>R</sup> | This study |
| pMET359 | 5,75 kb; pMET340 + *bla*; Erm<sup>R</sup>; Amp<sup>R</sup> | This study |
| pMET364 | pMET348::*attB*<sub>1</sub>; Cm<sup>R</sup> | This study |
| pMET365 | pMET348::*attB*<sub>3</sub>; Cm<sup>R</sup> | This study |
| pMET369 | pFSKB3X with 6xHisHUIIa under the control of lac promoter | This study |
| pMET370 | pMET348::*attB*<sub>2</sub>; Cm<sup>R</sup> | This study |
| pMET371 | pMET359::*core attP*<sub>1</sub>; Erm<sup>R</sup>, Amp<sup>R</sup> | This study |
| pMET372 | pMET359::*core attP*<sub>2</sub>; Erm<sup>R</sup>, Amp<sup>R</sup> | This study |
| pMET373 | pMET359::*core attP*<sub>3</sub>; Erm<sup>R</sup>, Amp<sup>R</sup> | This study |
| pMET376 | pMET359::*core attP*<sub>6</sub>; Erm<sup>R</sup>, Amp<sup>R</sup> | This study |

**Table S3.**
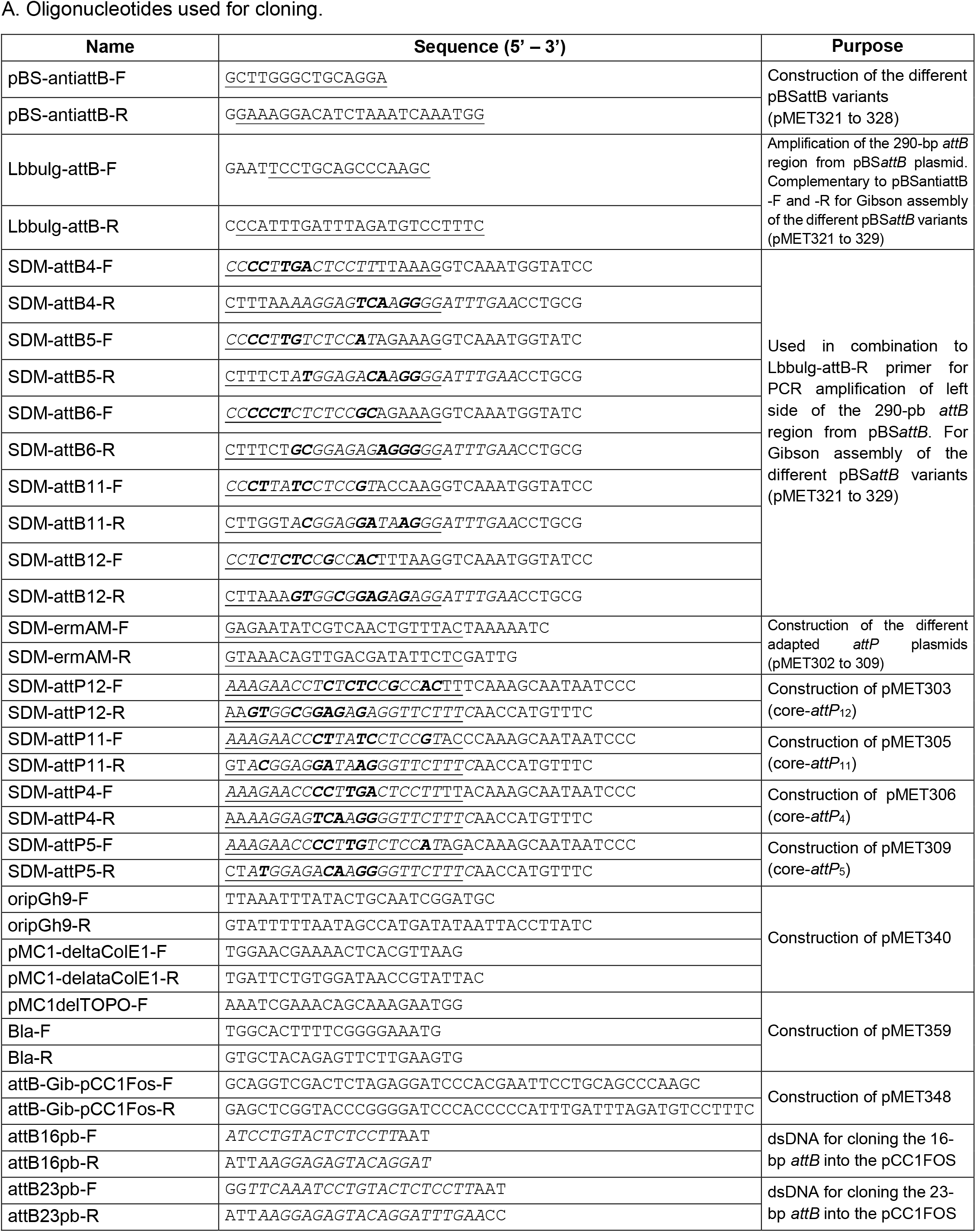
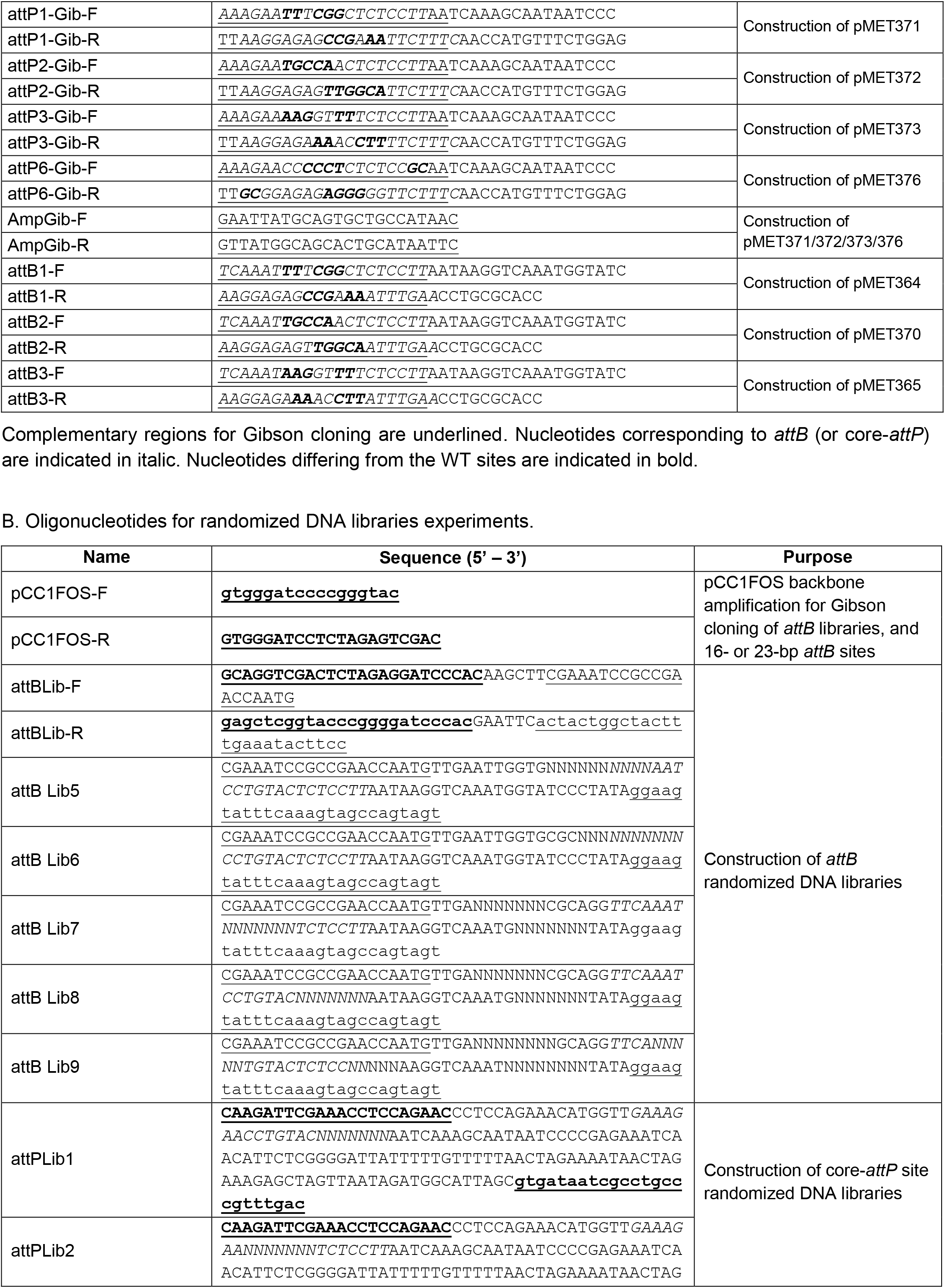
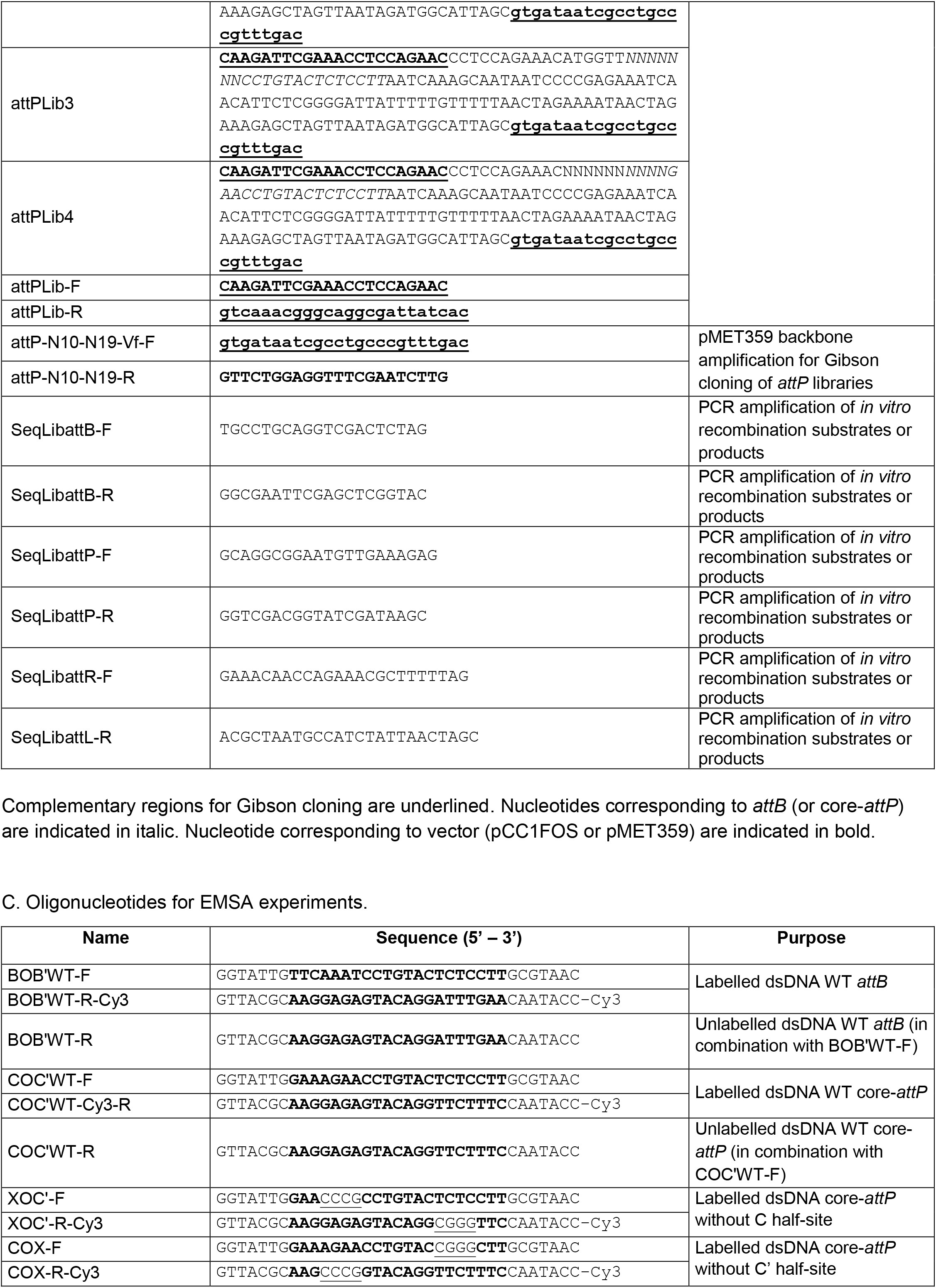
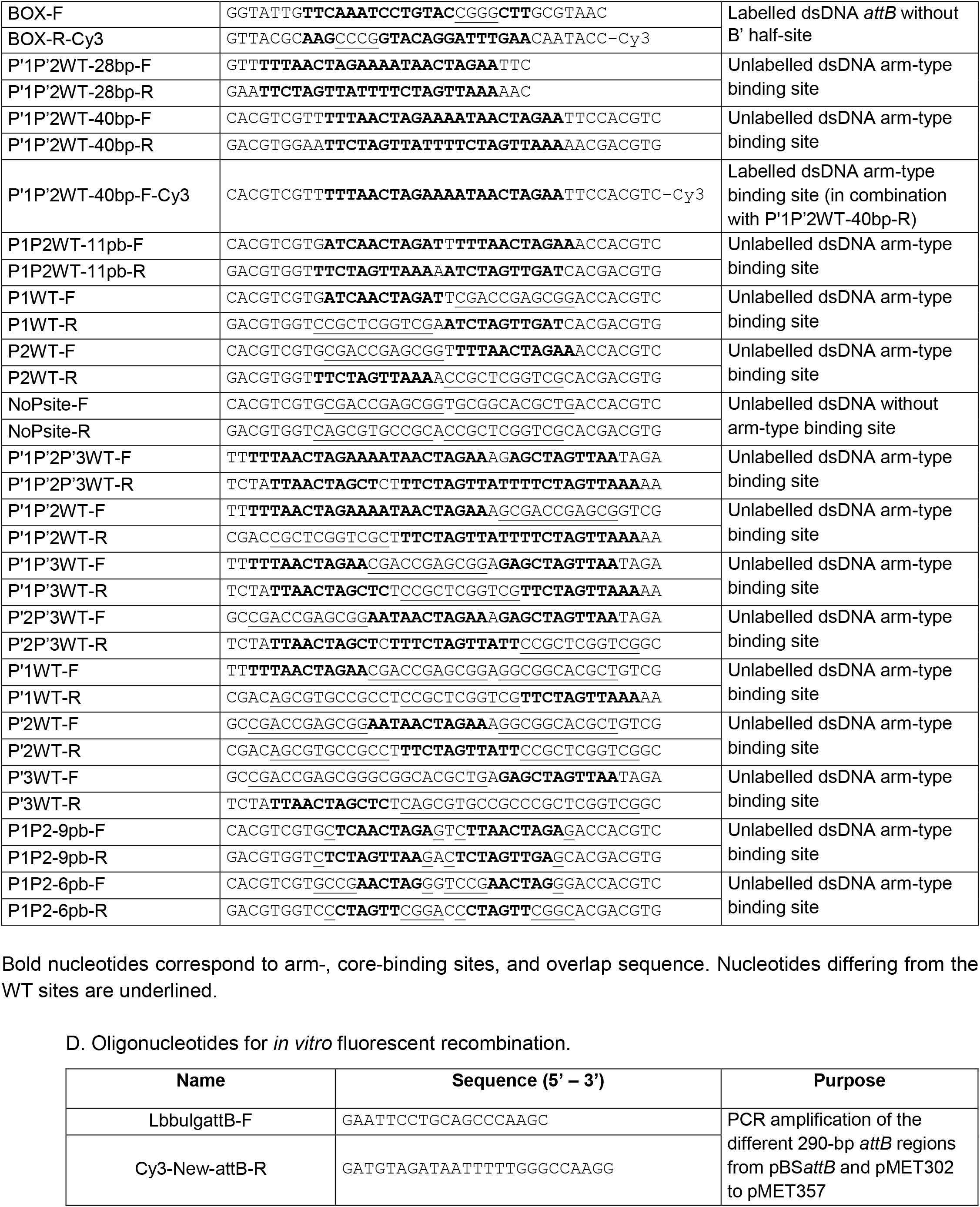
Oligonucleotide list.

**Table S4.** NGS data of the randomized DNA libraries.

| site | Reads after recombination <sup>a</sup> |  | Heptamers before recombination (*attP* or *attB*) | Heptamers after recombination |  | Heptamers present in both trials | Maximum read counts after recombination |  | Number of heptamers representing 50% of the total read count after recombination |  |
| --- | --- | --- | --- | --- | --- | --- | --- | --- | --- | --- |
|  | rep1 | rep2 |  | rep1 | rep2 |  | rep1 | rep2 | rep1 | rep2 |
| *attL* from *attB*Lib6 x *attP*<sub>WT</sub> (B) | 1 354 160 | 988 022 | 16 384 (100%)<sup>b</sup> | 5 459 (33,32%) | 6 955 (42,45%) | 4842 (33%) | 14 405 | 11 737 | 196 (3,59%) | 199 (2,86%) |
| *attR* from *attB*Lib8 x *attP*<sub>WT</sub> (B') | 1 814 444 | 2 352 851 | 16 312 (99,56%) | 9 497 (57,97%) | 10 085 (61,55%) | 8472 (52%) | 49 310 | 69 525 | 371 (3,91%) | 460 (4,56%) |
| *attR* from *attB*<sub>WT</sub> x *attP*Lib3 (C) | 1 434 050 | 2 754 031 | 16 340 (99,73%) | 10 416 (63,57%) | 11 781 (71,91%) | 9973 (61%) | 2 139 | 3 925 | 784 (7,53%) | 799 (6,78%) |
| *attL* from *attB*<sub>WT</sub> x *attP*Lib1 (C') | 930 830 | 1 447 157 | 16 315 (99,58%) | 7 549 (46,08%) | 7 247 (44,23%) | 5699 (35%) | 16 157 | 46 672 | 254 (3,36%) | 167 (2,30%) |

<sup>a</sup>: data from two independent trials.<sup>b</sup>:percentage of the theoretical heptamer population (4<sup>7</sup>) are indicate into brackets.

## REFERENCES

1. Casjens, S.R. and Hendrix, R.W. (2015) Bacteriophage lambda: early pioneer and still relevant. Virology, 479–480, 310–330.

2. Zeng, L., Skinner, S.O., Zong, C., Sippy, J., Feiss, M. and Golding, I. (2010) Decision making at a subcellular level determines the outcome of bacteriophage infection. Cell, 141, 682–691.

3. Silpe, J.E. and Bassler, B.L. (2019) A host-produced quorum-sensing autoinducer controls a phage lysis-lysogeny decision. Cell, 176, 268–280.e13.

4. Erez, Z., Steinberger-Levy, I., Shamir, M., Doron, S., Stokar-Avihail, A., Peleg, Y., Melamed, S., Leavitt, A., Savidor, A., Albeck, S., et al. (2017) Communication between viruses guides lysis-lysogeny decisions. Nature, 541, 488–493.

5. Grindley, N.D.F., Whiteson, K.L. and Rice, P.A. (2006) Mechanisms of site-specific recombination. Annu Rev Biochem, 75, 567–605.

6. Touchon, M., Bernheim, A. and Rocha, E.P. (2016) Genetic and life-history traits associated with the distribution of prophages in bacteria. ISME J, 10, 2744–2754.

7. Menouni, R., Hutinet, G., Petit, M.-A. and Ansaldi, M. (2015) Bacterial genome remodeling through bacteriophage recombination. FEMS Microbiol Lett, 362, 1–10.

8. Feiner, R., Argov, T., Rabinovich, L., Sigal, N., Borovok, I. and Herskovits, A.A. (2015) A new perspective on lysogeny: prophages as active regulatory switches of bacteria. Nat Rev Microbiol, 13, 641–650.

9. Casjens, S. (2003) Prophages and bacterial genomics: what have we learned so far? Mol Microbiol, 49, 277–300.

10. Gandon, S. (2016) Why Be Temperate: Lessons from Bacteriophage λ. Trends in Microbiology, 24, 356–365.

11. Rutherford, K. and Van Duyne, G.D. (2014) The ins and outs of serine integrase site-specific recombination. Curr Opin Struct Biol, 24, 125–131.

12. Lewis, J.A. and Hatfull, G.F. (2001) Control of directionality in integrase-mediated recombination: examination of recombination directionality factors (RDFs) including Xis and Cox proteins. Nucleic Acids Res, 29, 2205–2216.

13. Landy, A. (2015) The λ Integrase site-specific recombination pathway. Microbiol Spectr, 3, MDNA3-0051–2014.

14. de Vargas, L.M., Pargellis, C.A., Hasan, N.M., Bushman, E.W. and Landy, A. (1988) Autonomous DNA binding domains of λ integrase recognize two different sequence families. Cell, 54, 923–929.

15. Sarkar, D., Azaro, M.A., Aihara, H., Papagiannis, C.V., Tirumalai, R., Nunes-Düby, S.E., Johnson, R.C., Ellenberger, T. and Landy, A. (2002) Differential Affinity and Cooperativity Functions of the Amino-terminal 70 Residues of λ Integrase. J Mol Biol, 324, 775–789.

16. Tirumalai, R.S., Healey, E. and Landy, A. (1997) The catalytic domain of lambda site-specific recombinase. Proc Natl Acad Sci USA, 94, 6104–6109.

17. Gibb, B., Gupta, K., Ghosh, K., Sharp, R., Chen, J. and Van Duyne, G.D. (2010) Requirements for catalysis in the Cre recombinase active site. Nucleic Acids Res, 38, 5817–5832.

18. Better, M., Lu, C., Williams, R.C. and Echols, H. (1982) Site-specific DNA condensation and pairing mediated by the Int protein of bacteriophage lambda. Proc Natl Acad Sci USA, 79, 5837–5841.

19. Richet, E., Abcarian, P. and Nash, H.A. (1986) The interaction of recombination proteins with supercoiled DNA: Defining the role of supercoiling in lambda integrative recombination. Cell, 46, 1011–1021.

20. de Vargas, L.M., Kim, S. and Landy, A. (1989) DNA looping generated by DNA bending protein IHF and the two domains of lambda integrase. Science, 244, 1457–1461.

21. Richet, E., Abcarian, P. and Nash, H.A. (1988) Synapsis of attachment sites during lambda integrative recombination involves capture of a naked DNA by a protein-DNA complex. Cell, 52, 9–17.

22. Laxmikanthan, G., Xu, C., Brilot, A.F., Warren, D., Steele, L., Seah, N., Tong, W., Grigorieff, N., Landy, A. and Van Duyne, G.D. (2016) Structure of a Holliday junction complex reveals mechanisms governing a highly regulated DNA transaction. eLife, 5, e14313.

23. Seah, N.E., Warren, D., Tong, W., Laxmikanthan, G., Van Duyne, G.D. and Landy, A. (2014) Nucleoprotein architectures regulating the directionality of viral integration and excision. Proc Natl Acad Sci USA, 111, 12372–12377.

24. Tong, W., Warren, D., Seah, N.E., Laxmikanthan, G., Van Duyne, G.D. and Landy, A. (2014) Mapping the λ Integrase bridges in the nucleoprotein Holliday junction intermediates of viral integrative and excisive recombination. Proc Natl Acad Sci USA, 111, 12366–12371.

25. Smyshlyaev, G., Bateman, A. and Barabas, O. (2021) Sequence analysis of tyrosine recombinases allows annotation of mobile genetic elements in prokaryotic genomes. Mol Syst Biol, 17, e9880.

26. Esposito, D., Thrower, J.S. and Scocca, J.J. (2001) Protein and DNA requirements of the bacteriophage HP1 recombination system: a model for intasome formation. Nucleic Acids Res, 29, 3955–3964.

27. Panis, G., Méjean, V. and Ansaldi, M. (2007) Control and regulation of KplE1 prophage site-specific recombination: a new recombination module analyzed. J Biol Chem, 282, 21798–21809.

28. Sylwan, L., Frumerie, C. and Haggård-Ljungquist, E. (2010) Identification of bases required for P2 integrase core binding and recombination. Virology, 404, 240–245.

29. Smith-Mungo, L., Chan, I.T. and Landy, A. (1994) Structure of the P22 *att* site. Conservation and divergence in the lambda motif of recombinogenic complexes. J Biol Chem, 269, 20798–20805.

30. Lewis, J.A. and Hatfull, G.F. (2003) Control of directionality in L5 integrase-mediated site-specific recombination. J Mol Biol, 326, 805–821.

31. Lu, F. and Churchward, G. (1994) Conjugative transposition: Tn*916* integrase contains two independent DNA binding domains that recognize different DNA sequences. EMBO J, 13, 1541–1548.

32. Wood, M.M., DiChiara, J.M., Yoneji, S. and Gardner, J.F. (2010) CTnDOT integrase interactions with attachment site DNA and control of directionality of the recombination reaction. J Bacteriol, 192, 3934–3943.

33. Mattis, A.N., Gumport, R.I. and Gardner, J.F. (2008) Purification and characterization of bacteriophage P22 Xis protein. J Bacteriol, 190, 5781–5796.

34. Peña, C.E.A., Kahlenberg, J.M. and Hatfull, G.F. (2000) Assembly and activation of site-specific recombination complexes. Proc Natl Acad Sci USA, 97, 7760–7765.

35. Panis, G., Duverger, Y., Champ, S. and Ansaldi, M. (2010) Protein binding sites involved in the assembly of the KplE1 prophage intasome. Virology, 404, 41–50.

36. Cluzel, P.-J., Veaux, M., Rousseau, M. and Accolas, J.-P. (1987) Evidence for temperate bacteriophages in two strains of *Lactobacillus bulgaricus*. J Dairy Res, 54, 397–405.

37. Mata, M., Trautwetter, A., Luthaud, G. and Ritzenthaler, P. (1986) Thirteen virulent and temperate bacteriophages of *Lactobacillus bulgaricus* and *Lactobacillus lactis* belong to a single DNA homology group. Appl Environ Microbiol, 52, 812–818.

38. Dupont, L., Boizet-Bonhoure, B., Coddeville, M., Auvray, F. and Ritzenthaler, P. (1995) Characterization of genetic elements required for site-specific integration of *Lactobacillus delbrueckii* subsp. *bulgaricus* bacteriophage mv4 and construction of an integration-proficient vector for *Lactobacillus plantarum*. J Bacteriol, 177, 586–595.

39. Auvray, F., Coddeville, M., Espagno, G. and Ritzenthaler, P. (1999) Integrative recombination of *Lactobacillus delbrueckii* bacteriophage mv4: functional analysis of the reaction and structure of the *attP* site. Mol Gen Genet, 262, 355–366.

40. Auvray, F., Coddeville, M., Ordonez, R.C. and Ritzenthaler, P. (1999) Unusual structure of the *attB* site of the site-specific recombination system of *Lactobacillus delbrueckii* bacteriophage mv4. J Bacteriol, 181, 7385–7389.

41. Auvray, F., Coddeville, M., Ritzenthaler, P. and Dupont, L. (1997) Plasmid integration in a wide range of bacteria mediated by the integrase of *Lactobacillus delbrueckii* bacteriophage mv4. J Bacteriol, 179, 1837–1845.

42. Coddeville, M., Spinella, J.F., Cassart, P., Girault, G., Daveran-Mingot, M.L., Le Bourgeois, P. and Ritzenthaler, P. (2014) Bacteriophage mv4 site-specific recombination: the central role of the P2 ^mv4^Int-binding site. J. Virol., 88, 1839–1842.

43. Leenhouts, K., Buist, G., Bolhuis, A., ten Berge, A., Kiel, J., Mierau, I., Dabrowska, M., Venema, G. and Kok, J. (1996) A general system for generating unlabelled gene replacements in bacterial chromosomes. Mol Gen Genet, 253, 217–224.

44. Datsenko, K.A. and Wanner, B.L. (2000) One-step inactivation of chromosomal genes in *Escherichia coli* K-12 using PCR products. Proc Natl Acad Sci USA, 97, 6640–6645.

45. Bertani, G. (1951) Studies on lysogenesis. I. The mode of phage liberation by lysogenic *Escherichia coli*. J Bacteriol, 62, 293–300.

46. Terzaghi, B.E. and Sandine, W.E. (1975) Improved medium for lactic streptococci and their bacteriophages. Appl. Microbiol., 29, 807–813.

47. Gibson, D.G., Young, L., Chuang, R.-Y., Venter, J.C., Hutchison, C.A. and Smith, H.O. (2009) Enzymatic assembly of DNA molecules up to several hundred kilobases. Nat Methods, 6, 343– 345.

48. Afgan, E., Baker, D., Batut, B., van den Beek, M., Bouvier, D., Čech, M., Chilton, J., Clements, D., Coraor, N., Grüning, B.A., et al. (2018) The Galaxy platform for accessible, reproducible and collaborative biomedical analyses: 2018 update. Nucleic Acids Res, 46, W537–W544.

49. Langmead, B. and Salzberg, S.L. (2012) Fast gapped-read alignment with Bowtie 2. Nat Methods, 9, 357–359.

50. Schloss, P.D., Westcott, S.L., Ryabin, T., Hall, J.R., Hartmann, M., Hollister, E.B., Lesniewski, R.A., Oakley, B.B., Parks, D.H., Robinson, C.J., et al. (2009) Introducing mothur: open-source, platform-independent, community-supported software for describing and comparing microbial communities. Appl Environ Microbiol, 75, 7537–7541.

51. Rice, P., Longden, I. and Bleasby, A. (2000) EMBOSS: The European Molecular Biology Open Software Suite. Trends Genet, 16, 276–277.

52. Shannon, C.E. (1948) A mathematical theory of communication. The Bell System Technical Journal, 27, 379–423.

53. Cramér, H. (1946) Mathematical methods of statistics Princeton University Press.

54. Jumper, J., Evans, R., Pritzel, A., Green, T., Figurnov, M., Ronneberger, O., Tunyasuvunakool, K., Bates, R., Žídek, A., Potapenko, A., et al. (2021) Highly accurate protein structure prediction with AlphaFold. Nature, 596, 583–589.

55. Aihara, H., Kwon, H.J., Nunes-Düby, S.E., Landy, A. and Ellenberger, T. (2003) A conformational switch controls the DNA cleavage activity of lambda integrase. Mol Cell, 12, 187–198.

56. Wojciak, J.M., Sarkar, D., Landy, A. and Clubb, R.T. (2002) Arm-site binding by lambda -integrase: solution structure and functional characterization of its amino-terminal domain. Proc Natl Acad Sci USA, 99, 3434–3439.

57. Rubio-Cosials, A., Schulz, E.C., Lambertsen, L., Smyshlyaev, G., Rojas-Cordova, C., Forslund, K., Karaca, E., Bebel, A., Bork, P. and Barabas, O. (2018) Transposase-DNA complex structures reveal mechanisms for conjugative transposition of antibiotic resistance. Cell, 173, 208–220.e20.

58. Grove, A. (2011) Functional evolution of bacterial histone-like HU proteins. Curr Issues Mol Biol, 13, 1–12.

59. Sheren, J., Langer, S.J. and Leinwand, L.A. (2007) A randomized library approach to identifying functional *lox* site domains for the Cre recombinase. Nucleic Acids Res, 35, 5464–5473.

60. Malanowska, K., Salyers, A.A. and Gardner, J.F. (2006) Characterization of a conjugative transposon integrase, IntDOT. Mol Microbiol, 60, 1228–1240.

61. Craig, N.L. and Nash, H.A. (1983) The mechanism of phage lambda site-specific recombination: site-specific breakage of DNA by Int topoisomerase. Cell, 35, 795–803.

62. Kolot, M. and Yagil, E. (1994) Position and direction of strand exchange in bacteriophage HK022 integration. Mol Gen Genet, 245, 623–627.

63. Hauser, M.A. and Scocca, J.J. (1992) Site-specific integration of the *Haemophilus influenzae* bacteriophage HP1. Identification of the points of recombinational strand exchange and the limits of the host attachment site. J Biol Chem, 267, 6859–6864.

64. Peña, C.E., Stoner, J.E. and Hatfull, G.F. (1996) Positions of strand exchange in mycobacteriophage L5 integration and characterization of the *attB* site. J Bacteriol, 178, 5533–5536.

65. Hoess, R.H., Wierzbicki, A. and Abremski, K. (1986) The role of the *loxP* spacer region in P1 site-specific recombination. Nucleic Acids Res, 14, 2287–2300.

66. Weisberg, R.A., Enquist, L.W., Foeller, C. and Landy, A. (1983) Role for DNA homology in site-specific recombination. The isolation and characterization of a site affinity mutant of coliphage lambda. J Mol Biol, 170, 319–342.

67. Bauer, C.E., Gardner, J.F. and Gumport, R.I. (1985) Extent of sequence homology required for bacteriophage lambda site-specific recombination. J Mol Biol, 181, 187–197.

68. Kolot, M., Malchin, N., Elias, A., Gritsenko, N. and Yagil, E. (2015) Site promiscuity of coliphage HK022 integrase as a tool for gene therapy. Gene Ther, 22, 521–527.

69. McLeod, M., Craft, S. and Broach, J.R. (1986) Identification of the crossover site during FLP-mediated recombination in the *Saccharomyces cerevisiae* plasmid 2 microns circle. Mol Cell Biol, 6, 3357–3367.

70. Lee, G. and Saito, I. (1998) Role of nucleotide sequences of *loxP* spacer region in Cre-mediated recombination. Gene, 216, 55–65.

71. Missirlis, P.I., Smailus, D.E. and Holt, R.A. (2006) A high-throughput screen identifying sequence and promiscuity characteristics of the *loxP* spacer region in Cre-mediated recombination. BMC Genomics, 7, 1–13.

72. Nunes-Düby, S.E., Yu, D. and Landy, A. (1997) Sensing homology at the strand-swapping step in lambda excisive recombination. J Mol Biol, 272, 493–508.

73. Umlauf, S.W. and Cox, M.M. (1988) The functional significance of DNA sequence structure in a site-specific genetic recombination reaction. EMBO J, 7, 1845–1852.

74. Turan, S., Kuehle, J., Schambach, A., Baum, C. and Bode, J. (2010) Multiplexing RMCE: versatile extensions of the Flp-recombinase-mediated cassette-exchange technology. J Mol Biol, 402, 52–69.

75. Sarkar, D., Radman-Livaja, M. and Landy, A. (2001) The small DNA binding domain of lambda integrase is a context-sensitive modulator of recombinase functions. EMBO J, 20, 1203–1212.

76. Nolivos, S., Pages, C., Rousseau, P., Le Bourgeois, P. and Cornet, F. (2010) Are two better than one? Analysis of an FtsK/Xer recombination system that uses a single recombinase. Nucleic Acids Res, 38, 6477–6489.

77. Zhou, M., Bhasin, A. and Reznikoff, W.S. (1998) Molecular genetic analysis of transposase-end DNA sequence recognition: cooperativity of three adjacent base-pairs in specific interaction with a mutant Tn*5* transposase. J Mol Biol, 276, 913–925.

78. Cui, Z., Geurts, A.M., Liu, G., Kaufman, C.D. and Hackett, P.B. (2002) Structure-function analysis of the inverted terminal repeats of the sleeping beauty transposon. J Mol Biol, 318, 1221–1235.

79. Connolly, K.M., Iwahara, M. and Clubb, R.T. (2002) Xis protein binding to the left arm stimulates excision of conjugative transposon Tn*916*. J Bacteriol, 184, 2088–2099.

80. Johnson, R.C., Bruist, M.F. and Simon, M.I. (1986) Host protein requirements for in vitro site-specific DNA inversion. Cell, 46, 531–539.

81. Alonso, J.C., Weise, F. and Rojo, F. (1995) The Bacillus subtilis histone-like protein Hbsu is required for DNA resolution and DNA inversion mediated by the beta recombinase of plasmid pSM19035. J Biol Chem, 270, 2938–2945.

82. Petit, M.A., Ehrlich, D. and Jannière, L. (1995) pAM beta 1 resolvase has an atypical recombination site and requires a histone-like protein HU. Mol Microbiol, 18, 271–282.

83. Rowland, S.-J., Stark, W.M. and Boocock, M.R. (2002) Sin recombinase from *Staphylococcus aureus*: synaptic complex architecture and transposon targeting. Mol Microbiol, 44, 607–619.

84. Goodman, S.D. and Scocca, J.J. (1989) Nucleotide sequence and expression of the gene for the site-specific integration protein from bacteriophage HP1 of *Haemophilus influenzae*. J Bacteriol, 171, 4232–4240.

85. Yu, A. and Haggård-Ljungquist, E. (1993) Characterization of the binding sites of two proteins involved in the bacteriophage P2 site-specific recombination system. J Bacteriol, 175, 1239– 1249.

86. Pedulla, M.L., Lee, M.H., Lever, D.C. and Hatfull, G.F. (1996) A novel host factor for integration of mycobacteriophage L5. Proc Natl Acad Sci USA, 93, 15411–15416.

87. Ringwald, K. and Gardner, J. (2015) The *Bacteroides thetaiotaomicron* protein *Bacteroides* Host Factor A participates in integration of the integrative conjugative element CTnDOT into the chromosome. J Bacteriol, 197, 1339–1349.

88. Dey, D., Nagaraja, V. and Ramakumar, S. (2017) Structural and evolutionary analyses reveal determinants of DNA binding specificities of nucleoid-associated proteins HU and IHF. Mol Phylogenet Evol, 107, 356–366.

89. Kamashev, D., Agapova, Y., Rastorguev, S., Talyzina, A.A., Boyko, K.M., Korzhenevskiy, D.A., Vlaskina, A., Vasilov, R., Timofeev, V.I. and Rakitina, T.V. (2017) Comparison of histone-like HU protein DNA-binding properties and HU/IHF protein sequence alignment. PLoS One, 12, e0188037.

90. Swinger, K.K. and Rice, P.A. (2004) IHF and HU: flexible architects of bent DNA. Curr Opin Struct Biol, 14, 28–35.

91. Roberts, A.P. and Mullany, P. (2009) A modular master on the move: the Tn*916* family of mobile genetic elements. Trends Microbiol, 17, 251–258.

92. Alvarez, M.A., Herrero, M. and Suárez, J.E. (1998) The site-specific recombination system of the *Lactobacillus* species bacteriophage A2 integrates in gram-positive and gram-negative bacteria. Virology, 250, 185–193.

93. Raynal, A., Tuphile, K., Gerbaud, C., Luther, T., Guérineau, M. and Pernodet, J.-L. (1998) Structure of the chromosomal insertion site for pSAM2: functional analysis in *Escherichia coli*. Mol Microbiol, 28, 333–342.

94. Miele, S., Provan, J.I., Vergne, J., Possoz, C., Ochsenbein, F. and Barre, F.-X. (2022) The Xer activation factor of TLCΦ expands the possibilities for Xer recombination. Nucleic Acids Res, 50, 6368–6383.

95. Walker, M.W.G., Klompe, S.E., Zhang, D.J. and Sternberg, S.H. (2023) Novel molecular requirements for CRISPR RNA-guided transposition. Nucleic Acids Res, 51, 4519–4535.

96. Val, M.-E., Bouvier, M., Campos, J., Sherratt, D., Cornet, F., Mazel, D. and Barre, F.-X. (2005) The single-stranded genome of phage CTX Is the form used for integration into the genome of *Vibrio cholerae*. Mol Cell, 19, 559–566.

97. Das, B., Bischerour, J. and Barre, F.-X. (2011) VGJɸ integration and excision mechanisms contribute to the genetic diversity of *Vibrio cholerae* epidemic strains. Proc Natl Acad Sci USA, 108, 2516– 2521.

98. Bouvier, M., Demarre, G. and Mazel, D. (2005) Integron cassette insertion: a recombination process involving a folded single strand substrate. EMBO J, 24, 4356–4367.

99. Loot, C., Ducos-Galand, M., Escudero, J.A., Bouvier, M. and Mazel, D. (2012) Replicative resolution of integron cassette insertion. Nucleic Acids Res, 40, 8361–8370.

100. Rutkai, E., Dorgai, L., Sirot, R., Yagil, E. and Weisberg, R.A. (2003) Analysis of insertion into secondary attachment sites by phage lambda and by int mutants with altered recombination specificity. J Mol Biol, 329, 983–996.

101. Tanouchi, Y. and Covert, M.W. (2017) Combining Comprehensive Analysis of Off-Site Lambda Phage Integration with a CRISPR-Based Means of Characterizing Downstream Physiology. mBio, 8, e01038–17.

102. Elias, A., Kassis, H., Elkader, S.A., Gritsenko, N., Nahmad, A., Shir, H., Younis, L., Shannan, A., Aihara, H., Prag, G., et al. (2020) HK022 bacteriophage Integrase mediated RMCE as a potential tool for human gene therapy. Nucleic Acids Res, 48, 12804–12816.

103. Nunes-Duby, S.E., Kwon, H.J., Tirumalai, R.S., Ellenberger, T. and Landy, A. (1998) Similarities and differences among 105 members of the Int family of site-specific recombinases. Nucleic Acids Res., 26, 391–406.

104. Williams, K.P. (2002) Integration sites for genetic elements in prokaryotic tRNA and tmRNA genes: sublocation preference of integrase subfamilies. Nucleic Acids Res, 30, 866–875.

105. Biswas, T., Aihara, H., Radman-Livaja, M., Filman, D., Landy, A. and Ellenberger, T. (2005) A structural basis for allosteric control of DNA recombination by λ integrase. Nature, 435, 1059–1066.

## References

1. Aihara, H., Kwon, H.J., Nunes-Düby, S.E., Landy, A., and Ellenberger, T. (2003) A conformational switch controls the DNA cleavage activity of lambda integrase. Mol Cell 12: 187–198.

2. Auvray, F., Coddeville, M., Espagno, G., and Ritzenthaler, P. (1999a) Integrative recombination of *Lactobacillus delbrueckii* bacteriophage mv4: functional analysis of the reaction and structure of the *attP* site. Mol Gen Genet 262: 355–366.

3. Auvray, F., Coddeville, M., Ordonez, R.C., and Ritzenthaler, P. (1999b) Unusual structure of the *attB* site of the site-specific recombination system of *Lactobacillus delbrueckii* bacteriophage mv4. J Bacteriol 181: 7385–7389.

4. Coddeville, M., and Ritzenthaler, P. (2010) Control of directionality in bacteriophage mv4 site-specific recombination: functional analysis of the Xis factor. J Bacteriol 192: 624–635.

5. Coddeville, M., Spinella, J.F., Cassart, P., Girault, G., Daveran-Mingot, M.L., Le Bourgeois, P., and Ritzenthaler, P. (2014) Bacteriophage mv4 site-specific recombination: the central role of the P2 ^mv4^Int-binding site. J Virol 88: 1839–1842.

6. Corpet, F. (1988) Multiple sequence alignment with hierarchical clustering. Nucleic Acids Res 16: 10881– 10890.

7. Dupont, L., Boizet-Bonhoure, B., Coddeville, M., Auvray, F., and Ritzenthaler, P. (1995) Characterization of genetic elements required for site-specific integration of *Lactobacillus delbrueckii* subsp. *bulgaricus* bacteriophage mv4 and construction of an integration-proficient vector for *Lactobacillus plantarum*. J Bacteriol 177: 586–595.

8. Gibb, B., Gupta, K., Ghosh, K., Sharp, R., Chen, J., and Van Duyne, G.D. (2010) Requirements for catalysis in the Cre recombinase active site. Nucleic Acids Res 38: 5817–5832.

9. Smyshlyaev, G., Bateman, A., and Barabas, O. (2021) Sequence analysis of tyrosine recombinases allows annotation of mobile genetic elements in prokaryotic genomes. Mol Syst Biol 17: e9880.

10. Studier, F.W., Rosenberg, A.H., Dunn, J.J., and Dubendorff, J.W. (1990) Use of T7 RNA polymerase to direct expression of cloned genes. In Methods in Enzymology. Academic Press, pp. 60–89.

